# Lung cancer models reveal SARS-CoV-2-induced EMT contributes to COVID-19 pathophysiology

**DOI:** 10.1101/2020.05.28.122291

**Authors:** C. Allison Stewart, Carl M. Gay, Kavya Ramkumar, Kasey R. Cargill, Robert J. Cardnell, Monique B. Nilsson, Simon Heeke, Elizabeth M. Park, Samrat T. Kundu, Lixia Diao, Qi Wang, Li Shen, Yuanxin Xi, Bingnan Zhang, Carminia Maria Della Corte, Youhong Fan, Kiran Kundu, Boning Gao, Kimberley Avila, Curtis R. Pickering, Faye M. Johnson, Jianjun Zhang, Humam Kadara, John D. Minna, Don L. Gibbons, Jing Wang, John V. Heymach, Lauren Averett Byers

**Affiliations:** Department of Thoracic/Head & Neck Medical Oncology, The University of Texas MD Anderson Cancer Center, Houston, TX, USA; Department of Bioinformatics and Computational Biology, The University of Texas MD Anderson Cancer Center, Houston, TX, USA; Department of Precision Medicine, Oncology Division, University of Campania “Luigi Vanvitelli”, Naples, Italy; Department of Internal Medicine and Pharmacology, Hamon Center for Therapeutic Oncology Research, The University of Texas Southwestern Medical Center, Dallas, TX, USA; Department of Head and Neck Surgery, The University of Texas MD Anderson Cancer Center, Houston, TX, USA; Department of Translational Molecular Pathology, The University of Texas MD Anderson Cancer Center, Houston, TX, USA

**Author notes:** These authors contributed equally to this work (C. Allison Stewart, Carl M. Gay).

## Abstract

COVID-19 is an infectious disease caused by SARS-CoV-2, which enters host cells via the cell surface proteins ACE2 and TMPRSS2. Using a variety of normal and malignant models and tissues from the aerodigestive and respiratory tracts, we investigated the expression and regulation of *ACE2* and *TMPRSS2*. We find that *ACE2* expression is restricted to a select population of highly epithelial cells. Notably, infection with SARS-CoV-2 in cancer cell lines, bronchial organoids, and patient nasal epithelium, induces metabolic and transcriptional changes consistent with epithelial to mesenchymal transition (EMT), including upregulation of *ZEB1* and *AXL*, resulting in an increased EMT score. Additionally, a transcriptional loss of genes associated with tight junction function occurs with SARS-CoV-2 infection. The SARS-CoV-2 receptor, ACE2, is repressed by EMT via TGFbeta, ZEB1 overexpression and onset of EGFR TKI inhibitor resistance. This suggests a novel model of SARS-CoV-2 pathogenesis in which infected cells shift toward an increasingly mesenchymal state, associated with a loss of tight junction components with acute respiratory distress syndrome-protective effects. AXL-inhibition and ZEB1-reduction, as with bemcentinib, offers a potential strategy to reverse this effect. These observations highlight the utility of aerodigestive and, especially, lung cancer model systems in exploring the pathogenesis of SARS-CoV-2 and other respiratory viruses, and offer important insights into the potential mechanisms underlying the morbidity and mortality of COVID-19 in healthy patients and cancer patients alike.

## Introduction

In December 2019, reports of a viral illness with severe respiratory symptoms emerged from Wuhan, China^1,2^. The novel virus was classified as a coronavirus and subsequently named SARS-CoV-2. It exhibited rapid global spread and met the World Health Organization’s criteria of a pandemic within three months following the first reported diagnosis (who.int). As of January 27, 2021, more than 99 million individuals have been infected worldwide with over 2.1 million deaths (covid19.who.int). SARS-CoV-2 infection is the cause of the respiratory illness COVID-19, which presents most frequently with symptoms including cough and dyspnea, accompanied by a moderate to high fever^2,3^. The severity of patient symptoms following SARS-CoV-2 infection varies widely from asymptomatic carrier-status to critical illness. Given the specific complications associated with infection, it is not initially surprising that patients with cancer and, specifically, thoracic malignancies seem to experience poorer clinical outcomes^4,5^. However, it is unclear whether these poorer outcomes represent the impact of demographics (e.g. age, smoking status, gender) or cellular and molecular changes associated with the tumor and its microenvironment.

While there are currently no validated molecular biomarkers for susceptibility to, or severity of, SARS-CoV-2 infection, it has recently been described that, as with SARS-CoV, SARS-CoV-2 cell entry requires interactions with angiotensin-converting enzyme 2 (ACE2) and transmembrane serine protease 2 (TMPRSS2) on the surface of the host cell^6^. Specifically, ACE2 binds to a subunit of the SARS-CoV-2 spike (S) protein, while TMPRSS2 is responsible for S-protein cleavage, or priming, to allow fusion of viral and host cellular membranes^6^. Differences in these receptors has been described in certain populations harboring higher allelic frequency of mutations and coding variants associated with higher *ACE2* expression^7^. Notably, ACE2 has previously been shown to exert a protective effect toward the development of acute respiratory distress syndrome (ARDS) – one of the most common and lethal complications of SARS-CoV-2 infection^8,9^. Given the propensity for ARDS in SARS-CoV-2 infected patients, this protective effect initially seems paradoxical, but previous data on SARS-CoV-1 suggest that subsequent to infection ACE2 expression is downregulated, thus tipping the balance in favor of acute lung injury^10^.

As the presence of ACE2 and/or TMPRSS2 may be rate-limiting for SARS-CoV-2 infection, we utilized bulk and single-cell transcriptional data from a combination of normal and malignant tissue samples and models from the aerodigestive and respiratory tracts to explore mechanisms governing the expression of *ACE2* and *TMPRSS2*. Our bulk data suggests that aerodigestive and lung cancer cell lines have a broad range of *ACE2* and *TMRPSS2* expression and would serve as good models for studying SARS-CoV-2 infection, consistent with the observation that SARS-CoV-2 virus has the ability to infect several lung cancer cell lines (e.g., Calu-3 and A549)^6,11^. Furthermore, while the *normal* aerodigestive and respiratory tracts represent key points of viral entry and infection, limiting an investigation to *normal* tissue may unnecessarily limit available samples for analysis and, especially, limit the range of cellular and molecular phenomena represented by those samples.

Meanwhile, single-cell analyses demonstrate that while *TMPRSS2* is widely expressed in normal respiratory epithelium, *ACE2* expression is limited to a small collection of cells^12^ and, therefore, may be a limiting factor for infection or drive dependence on other mechanisms, such as putative AXL-mediated cell entry^13-17^. There have been numerous reports of patients developing loss of chemosensation (taste, smell, etc.) following SARS-CoV-2 infection – an observation that pointed to another small collection of cells^18,19^. Tuft, or brush, cells are rare chemosensory cells present in the aerodigestive and respiratory tracts, among other sites, that mediate taste and smell, along with innate immune response^20,21^. These cells, which are epithelial in nature, strongly express *POU2F3*, a transcription factor that is also highly expressed in certain lung cancers, including a subset of small cell lung cancer^22-25^. Our analyses indeed suggest that *ACE2* expression is enriched in tuft cells compared to non-tuft cells within the respiratory tract. However, this was not absolute and, thus, we considered broader regulatory mechanisms that might govern *ACE2* expression. Previous literature has proposed, inconsistently, both positive and negative associations between *ACE2* expression and epithelial differentiation^26,27^, while tuft cells, in which *ACE2* is enriched in our analysis, are highly epithelial^21^. We assessed the relationship between *ACE2* and epithelial differentiation in numerous aerodigestive and respiratory datasets, and found striking and consistent positive correlations with transcriptional, microRNA, and metabolic signifiers of epithelial differentiation in virtually all datasets analyzed.

Finally, we consider the role that regulators of epithelial to mesenchymal transition (EMT) may play in modulation of *ACE2* expression. EMT is a well-characterized phenomenon critical to normal developmental processes, but co-opted by tumor cells to promote resistance and metastasis^28^. We provide evidence that the miR-200 family – zinc finger E-box-binding homeobox 1 (ZEB1) pathway, which is an established regulator of EMT^29,30^, also directly regulates *ACE2* expression, likely via putative ZEB1 repressor sites located in the *ACE2* promoter. Furthermore, we highlight that SARS-CoV-2 infection both *in vitro* and in patients yields increased expression of EMT-associated genes, including *ZEB1* and *AXL* and metabolic derangements (e.g. a shift away from glutamine and toward glutamate) characteristic of EMT^31,32^. Inhibition of AXL, a mesenchymal receptor tyrosine kinase, with bemcentinib both induces ACE2 and represses ZEB1 and may serve as a therapy to shift SARS-CoV-2 infected cells away from a mesenchymal phenotype (i.e., reverse EMT). Additionally, viral infection of epithelial cells, which are glutamine-dependent with high *GLUL* expression and presumably more metabolically-primed for replication, promotes rapid replication due to the enhanced dependence on glutamine for nucleotide synthesis and subsequently depletes glutamine stores^32^.

The association between EMT and lung injury/ARDS has been described previously, with the central pathogenesis being the breakdown of the alveolar epithelial barrier^33,34^. Intriguingly, previous data demonstrated that over-expression of miR-200 family members (which inhibit *ZEB1* expression) or direct exogenous silencing of ZEB1 has a protective effect in a murine, lipopolysaccharide-induced model of ARDS^35^. In this context, our findings support a novel model for the pathogenesis of SARS-CoV-2 in which the virus initially infects a small pool of highly epithelial cells of the aerodigestive and respiratory tracts followed by those infected cells undergoing molecular alterations typical of EMT (i.e., *ZEB1/AXL* upregulation). These EMT-like alterations, in turn, result directly in the down-regulation of genes associated with tight junctions and, in doing so, eradicate the proposed ARDS-protective effect of these epithelial, ACE2-positive cells.

## Results

### Expression of ACE2 by cancer epithelial cells

*ACE2* is widely expressed in normal tissues from Genotype-Tissue Expression (GTEx) Project **(http://www.gtexportal.org)** and was highest in testis and small intestine and lowest in spleen and blood with moderate expression in lung and salivary glands, which are sites of SARS-CoV-2 transmission and/or infection (Supplemental Figure 1a). Transcriptional data is shown as normalized relative gene expression. High levels of *ACE2* in the small intestine may explain reports of gastrointestinal distress in COVID-19 patients^36^. To leverage cancer cell lines and patient tumors to SARS-CoV-2 research, we explored expression of the receptors, *ACE2* and *TMPRSS2*, and discovered a negative correlation with EMT. This was validated in SARS-CoV-2-infected samples and led to investigation of ways to manipulate EMT in an effort to minimize infection and/or ARDS in patients (Figure 1a). *ACE2* expression in aerodigestive and respiratory cancer cell lines (Figure 1b) and both normal and tumor specimens (Figure 1c) is consistently higher in specimens with a low EMT score, suggesting that *ACE2* is primarily expressed by epithelial cells in these cancers^37-48^. The EMT score uses 76 genes to define the degree to which cells or tissues have undergone EMT and has been applied in TCGA projects^49-51^. This observation is confirmed by a consistent negative correlation of *ACE2* and EMT score in all nine cell line and tumor data sets (Table 1). *TMPRSS2*, the protease also required for SARS-CoV-2 infection, is similarly expressed in models with epithelial gene signatures (Figure 1b,c) and highly correlated with *ACE2* expression in cancer models (Table 1). To determine whether expression levels of *ACE2* were different in normal tissues and tumors, we compared expression in TCGA tumor datasets with paired normal, adjacent samples. Both normal and lung adenocarcinoma (LUAD) tissues have similar *ACE2* expression (TCGA LUAD P=0.21) and EMT score (TCGA LUAD P=0.87) (Supplemental Figure 1b), although a greater range of expression (both higher and lower) could be observed in tumors^52-54^. Interestingly, and in contrast to some previous reports, no difference in *ACE2* expression was detected in normal lung tissue (adjacent to TCGA LUAD) with regards to smoking status, gender or age (Supplemental Figure 1c).

**Figure 1.**
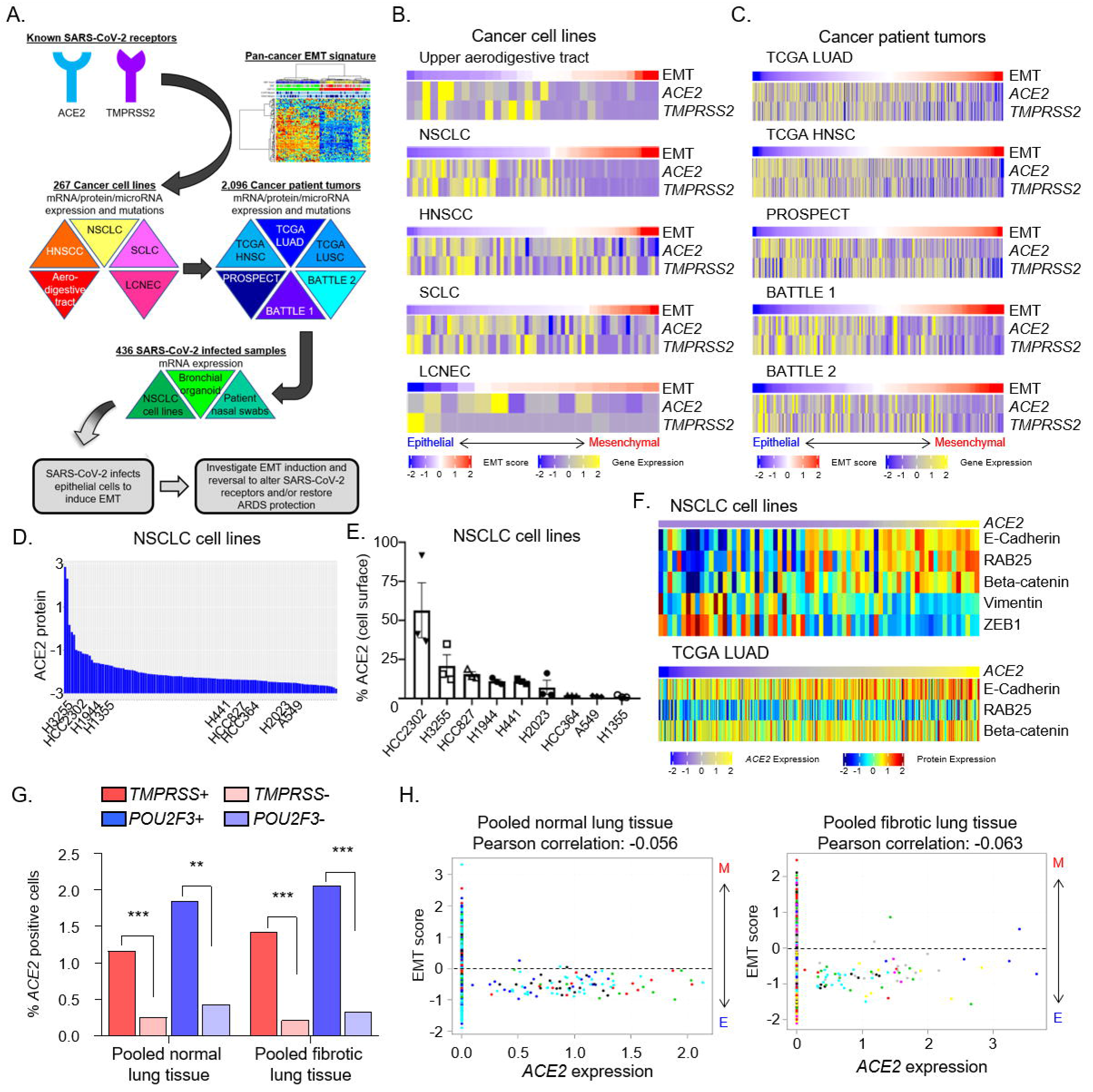
The SARS-CoV2 receptor, *ACE2*, is expressed by epithelial cells in cancer models. a, Schematic depicting an overview of the strategy to investigate SARS-CoV-2 receptor biology in the context of EMT using cancer cell lines and tumor samples before applying to SARS-CoV-2-infected samples to develop our hypotheses. b, *ACE2* and *TMPRSS2* expression and EMT score in aerodigestive and lung cancer cell lines (*ACE2* vs. EMT score, P<0.0001 for all) and c, in patient tumor biopsies (*ACE2* vs, EMT score, P<0.0001 for all). Rho values are listed in Table 1. d, ACE2 protein levels in NSCLC cell lines. e, Cell surface ACE2 in a subset of NSCLC cell lines. f, NSCLC cell lines and TCGA LUAD tumor biopsies were ranked by *ACE2* mRNA expression and demonstrate protein expression of common epithelial (E-cadherin, RAB25, beta-catenin) or mesenchymal (vimentin, ZEB1, fibronectin) markers. g, Frequency of *ACE2*-positive cells in *TMPRSS2*-positive or -negative cells and *POU2F3*-positive or negative cells as determined by single-cell RNAseq of donor or fibrotic lungs. h, EMT score values for each *ACE2*-positive cell present in five donor and eight fibrotic lungs. Each color represents cells from an individual lung sample.

**Table 1.** ACE2 rho correlation values in aerodigestive and respiratory tract cell line and tumor biopsy specimens. *P<0.05, **P<0.01, ***P<0.001.

| Dataset | ACE2 expression correlated with: |  |  |  |  |  |  |
| --- | --- | --- | --- | --- | --- | --- | --- |
|  | *TMPRSS2* | *POU2F3* | EMT score | *ZEB1* | *AXL* | *GLUL* | *GLS* |
| **NSCLC cell lines** | 0.565*** | 0.668*** | -0.694*** | -0.801*** | -0.410*** | 0.394*** | -0.202* |
| **HNSCC cell lines** | 0.230* | 0.188 | -0.408*** | -0.388*** | -0.078 | 0.306** | -0.042 |
| **SCLC cell lines** | 0.069 | 0.394** | -0.401*** | -0.429*** | -0.351* | 0.284* | -0.305* |
| **TCGA LUAD** | 0.228*** | 0.328*** | -0.286*** | -0.28*** | -0.083 | 0.296*** | -0.197*** |
| **TCGA HNSC** | 0.082 | 0.159*** | -0.402*** | 0.238*** | -0.239*** | 0.037 | -0.071 |
| **TCGA LUSC** | 0.242*** | 0.033 | -0.239*** | -0.070 | -0.121** | 0.404*** | -0.179*** |
| **PROSPECT** | 0.195*** | 0.065 | -0.279*** | -0.117* | -0.090** | -0.091 | 0.126* |
| **BATTLE 1** | 0.437*** | 0.134 | -0.367*** | -0.166* | -0.222** | 0.339*** | -0.082 |
| **BATTLE 2** | 0.373*** | 0.249** | -0.337*** | -0.196* | -0.235** | 0.208** | -0.064 |

biopsy specimens. \*P<0.05, \*\*P<0.01, \*\*\*P<0.001.

In addition to *ACE2* mRNA expression, we screened a cohort of NSCLC and SCLC cell lines for ACE2 protein levels by reverse phase protein array (RPPA; Figure 1d) and within a subset of NSCLC cell lines we analyzed cell surface expression of NSCLC cell lines by flow cytometry (Figure 1e). Similar to gene expression, ACE2 levels negatively correlate with mesenchymal proteins, including vimentin (Supplemental Figure 1d). To go beyond the quantitative, gene-based EMT score, we investigated expression of *ACE2* mRNA compared to specific epithelial (i.e., E-cadherin, RAB25, beta-catenin) or mesenchymal (i.e., vimentin, ZEB1) proteins by RPPA analysis. *ACE2* was higher in non-small cell lung cancer (NSCLC) cell lines (P<0.008) and TCGA tumors (P<0.001; Figure 1f) with higher levels of epithelial proteins, consistent with the mRNA associations and EMT score.

*ACE2* is only expressed by a small subset of cells (0-4.7%) at the single-cell transcriptional level in normal respiratory tissues, as well as in oral cavity tumors and lung cancer xenograft models (Table 2)^55-59^. However, in all datasets, the rare, *ACE2*-positive cells consistently have a low, or more epithelial, EMT score. In both fibrotic and donor lung samples, *TMPRSS2* is more widely expressed than *ACE2*, while *ACE2* expression is found more frequently in *TMPRSS2*-positive cells (Figure 1g, Supplemental Table 1) relative to *TMPRSS2*-negative cells. As studies suggest both are required for infection, these data, indicate that *ACE2* expression is more limiting than *TMPRSS2* expression for SARS-CoV-2 infection within respiratory epithelium.

**Table 2.** Total number of *ACE2*-positive cells and EMT score.

| **Datasets** | **Total # of cells/samples** | ***ACE2*+ cells (%)** | **EMT score in *ACE2*+ cells** |
| --- | --- | --- | --- |
| Bronchial brushings (GSE131391) | 1146 cells (12 patients) | 54 (4.71%) | -0.13 |
| Oral cavity tumors (GSE103322) | 5902 cells (18 tumors) | 87 (1.47%) | -0.54 |
| Donor lungs (GSE122960) | 20,000 cells (5 lungs) | 94 (0.47%) | -0.52 |
| Fibrotic lungs (GSE122960) | 19,587 cells (8 lungs) | 72 (0.37%) | -0.75 |
| Parenchymal lung (GSE130148) | 10,360 cells (3 lungs) | 14 (0.14%) | -0.26 |
| PC9 xenograft DMSO treated (GSE138693) | 3766 cells (1 xenograft) | 29 (0.77%) | -0.40 |
| PC9 xenograft osimertinib treated (GSE138693) | 2,332 cells (1 xenograft) | 0 (0%) | N/A |
| Bronchial biopsy/Nasal brushing/Turbinate (GSE121600) | 12,000 cells (3 specimens) | 4 (0.03%) | -0.49 |

*POU2F3* is a marker of tuft cell expression and is correlated with *ACE2* expression in many aerodigestive and lung cancer data sets (Table 1). ACE2 protein levels also correlate with POU2F3 in both NSCLC (Rho=0.221, P=0.01) and SCLC (Rho=0.673, P<0.0001) cell lines. Similar to the observation for co-expression with *TMPRSS2, ACE2* expression is more often detected in *POU2F3*-positive than negative cells (Figure 1g, Supplemental Table 1). This suggests that tuft cells may be preferentially targeted by SARS-CoV-2 infection in aerodigestive tissues. These rare, *ACE2*-positive cells can be visualized in tSNE space (Supplemental Figure 2a) and demonstrate a low EMT score on a cell-by-cell basis in both donor and fibrotic lung samples (Figure 1h) and oral cavity tumors (Supplemental Figure 2b). These *ACE2*-positive cells are localized within epithelial cell clusters with low EMT score that primarily consist of alveolar type II, ciliated and club cells (Supplemental Figure 2c) based on expression of cell-type specific markers including (*SFTPC* [alveolar type II cells], *FOXJI* [ciliated cells], *SCGB1A1* [club cells]).

### SARS-CoV-2 infection induces EMT

We next sought to determine the impact of SARS-CoV-2 infection on the EMT status of the infected cell. We analyzed RNAseq data from NSCLC cell lines (A549 and Calu-3) infected for 24h with SARS-CoV-2 (GSE147507)^60^. Since A549 cells have low endogenous levels of *ACE2* expression, the authors additionally transduced these cells with a vector over-expressing human ACE2 to improve infection. Expression of the epithelial gene *EPCAM* was downregulated following viral infection (Calu-3 P=0.03, A549+ACE2 P=0.01 Figure 2a; A549 P=0.04 Supplemental Figure 3a), suggesting that infection is inducing a shift away from an epithelial phenotype. This was confirmed by *ZEB1* upregulation in all three cell lines (Calu-3 P=0.01, A549+ACE2 P=0.05 Figure 2a; A549 P=0.003 Supplemental Figure 3a) following infection.

**Figure 2.**
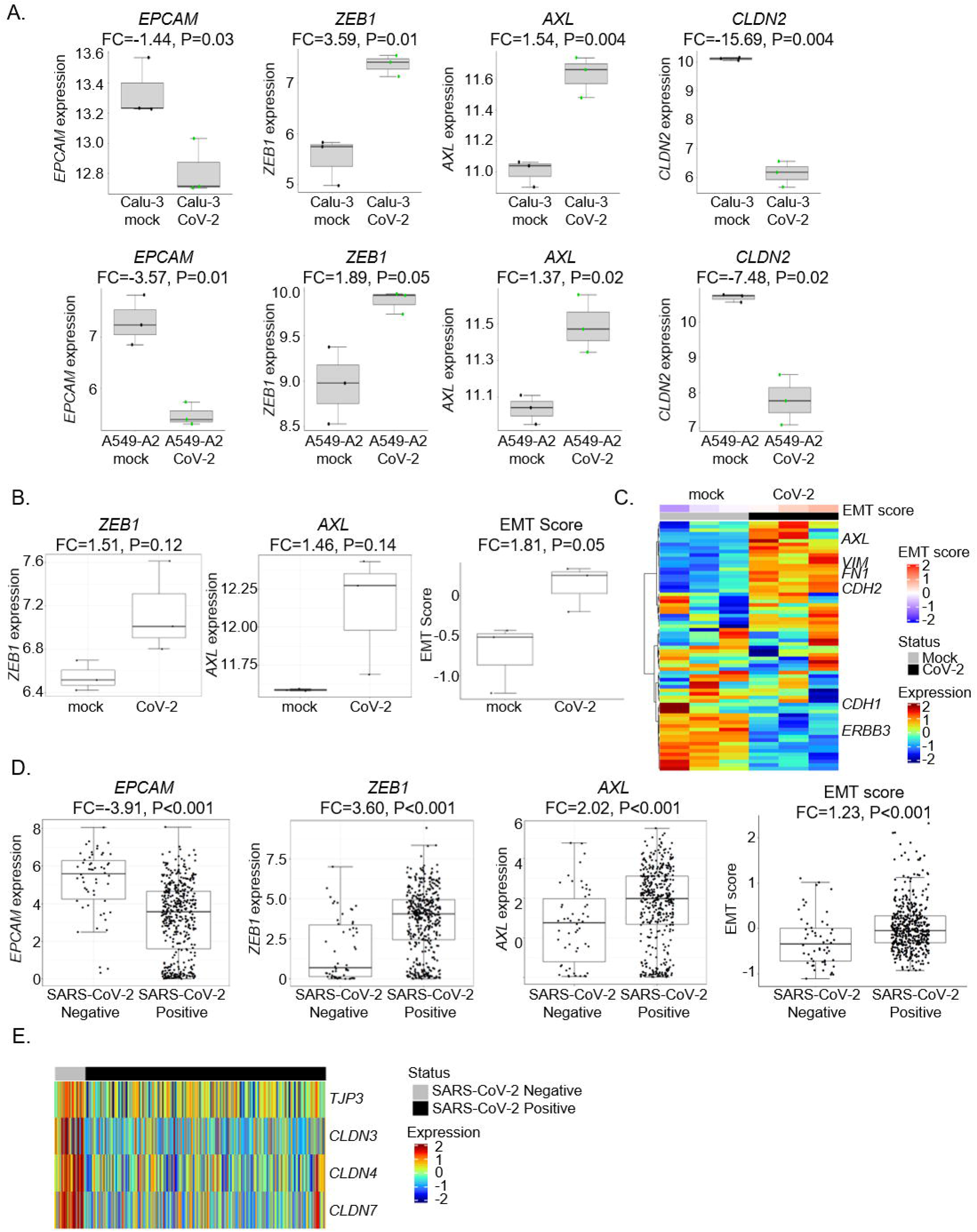
SARS-CoV-2 induces EMT. a, Effects of SARS-CoV-2 infection of Calu-3 or A549+ACE2 for 24h on *EPCAM, ZEB1, AXL*, and *CLDN2* expression. b, Effects of SARS-CoV-2 infection on human bronchial organoids for 5 days on *ZEB1* and *AXL* expression and EMT score. c, Heatmap demonstrating differences in gene expression between mock and SARS-CoV-2 infected bronchial organoids in EMT-associated genes. d, Effects of SARS-CoV-2 infection in nasal swab samples from 430 individuals with PCR-confirmed SARS-CoV-2 and 54 negative controls on *EPCAM, ZEB1, AXL* expression and EMT score. e, Effects of SARS-CoV-2 infection in nasal swabs on genes associated with tight junctions. RNAseq data is shown as normalized relative gene expression.

An alternative EMT regulator, AXL, which is a TAM (Tyro3, AXL, Mer) family receptor tyrosine kinase strongly associated with a mesenchymal phenotype, has emerged as a key determinant of therapeutic resistance in NSCLC and other cancer types^49,61^. Similar to *ZEB1, AXL* is inversely correlated with *ACE2* in cell line and tumor samples (Table 1). Furthermore, infection of cells with SARS-CoV-2 upregulated *AXL* expression in all three cell lines (Calu-3 P=0.004, A549+ACE2 P=0.02 Figure 2a; A549 P<0.001 Supplemental Figure 3a). A similar induction of mesenchymal genes with viral infection was observed for *SNAI1* (Calu-3: FC=1.84, P=0.009; A549+ACE2: FC=7.43, P=0.01) and *ZEB2* (Calu-3: FC=6.74, P=0.002; A549+ACE2: FC=5.17, P=0.0001). However, EMT score was not different between mock or SARS-CoV-2 infected Calu-3 (FC= −1.09, P=0.14) or A549+ACE2 cells (FC= −1.03, P=0.48) and this may be due to the relatively short period of viral infection (24h).

EMT is also known to downregulate tight junction components, including several members of the Claudin protein family^62^. Claudins are integral membrane proteins localized at tight junctions that are involved in regulating epithelial cell polarity and paracellular permeability. COVID-19 patients with ARDS suffer from pulmonary edema due in part to disruption of tight junctions within alveolar-epithelial barrier^63^. *CLDN2* is expressed in respiratory epithelium, but a role in mediating alveolar edema has not been described^64^. However, out of all Claudin family members, *CLDN2* was downregulated in all three cell lines (Calu-3 P=0.004, A549+ACE2 P=0.02 Figure 2a; A549 P=0.008 Supplemental Figure 3a), suggesting a change in epithelial and endothelial cell permeability that may contribute to ARDS in patients with COVID-19.

Similarly, there was a reduction in *TJP3*, the gene encoding a scaffolding protein that links tight junction transmembrane proteins, such as claudins, to the actin cytoskeleton (Calu3: FC= −1.48, P=0.04; A549+ACE2: FC=-2.89, P=0.10) following viral infection. Evidence of a metabolic shift away from glutamine was observed post-infection, as *GLUL* expression was reduced in *ACE2-*high cell lines, but not in non-transfected A549 (Calu-3 P=0.01, A549+ACE2 P=0.01 Supplemental Figure 3b, A549 P=0.56). The *ZEB1* and *AXL* increase and *EPCAM* and *GLUL* decrease point toward a SARS-CoV-2 induced shift from an epithelial to mesenchymal phenotype.

To determine whether longer exposure to virus may induce additional EMT changes, we analyzed RNAseq data from human bronchial organoids developed from commercially available, cryopreserved adult bronchial epithelial cells infected with SARS-CoV-2 for five days (GSE150819)^65^. There was a subtle increase in both *ZEB1* (P=0.12) and *AXL* (P=0.14), as well as other mesenchymal genes, including *VIM* (FC=1.56, P=0.01), *ZEB2* (FC=3.23, P=0.22), *SNAI1* (FC=4.75, P=0.05), *CDH2* (FC=2.06, P=0.02). Importantly, longer SARS-CoV-2 infection increased the EMT score (P=0.05; Figure 2b). While these shifts are not as strong as those seen in the cell lines, there is a clear pattern for expression of the epithelial and mesenchymal genes making up the EMT score by heatmap (Figure 2c). Similar to the NSCLC cell lines, SARS-CoV-2 infection downregulated expression of tight junction-related genes, including *CLDN8* (FC=-2.66, P=0.009) and *CLDN17* (FC=-1.62; P=0.05).

Finally, to evaluate whether viral-induced EMT also occurs in patients, we analyzed metagenomic next-generation sequencing data of human nasopharangeal swabs from 430 PCR-confirmed SARS-CoV-2 and 54 negative control individuals (GSE154770)^66^. SARS-CoV-2 infection downregulated expression of EPCAM (P<0.0001) and upregulated *ZEB1* (P<0.0001) and *AXL* (P<0.0005; Figure 2d). Similar patterns were found with other epithelial genes, including *RAB25* (FC=-2.35; P<0.0001) and *CTNNB1* (FC=-3.38; P<0.0001) or mesenchymal genes, including *ZEB2* (FC=8.14, P<0.0001). As expected, viral infection was associated with an increased EMT score (P<0.0001; Figure 2d) and *TGFB1* expression (FC=2.75, P<0.0001).

Interestingly, EMT gene expression was not impacted by viral load (Supplemental Figure 3c), as defined by the cycle threshold (Ct) of the SARS-CoV-2 nucleocapsid gene region 1 (N1) target during diagnostic PCR^66^. This suggests that EMT is induced in cells, regardless of the viral burden in the patient. Consistent with the previous metabolic data, SARS-CoV-2 infection induces expression of *GLS*, indicating a shift away from glutamine production (P=0.009; Supplemental Figure 3d). Similar to the cell lines and bronchial organoids, SARS-CoV-2 infection reduces expression of genes associated with tight junctions (Figure 2e), including *CLDN4* (FC=-6.63, P<0.0001), *CLDN7* (FC=-6.59, P<0.0001), *CLDN3* (FC=-5.14; P<0.0001), *CLDN9* (FC=-1.72; P<0.0001), *CLDN23* (FC=-1.95, P=0.0002), *TJP2* (FC=-1.58, P=0.05), and *TJP3* (FC=-2.22, P<0.0001).

### Induction of EMT represses ACE2 expression

To connect differences in *ACE2* expression to a more specific EMT regulatory program, we investigated miRNAs in both lung cancer and head and neck cell lines and tumor samples. Interestingly, miRNAs from the miR-200 family were consistently positively correlated with *ACE2*^*67*^. As expected, given their known role as inhibitors of EMT, high miR-200 family expression was seen in cells and tumors with high *CDH1* (an epithelial-specific marker) and low EMT score and *ZEB1*, the latter a transcriptional promoter of EMT and target of miR-200 family (Figure 3a; Supplemental Figure 3e). The miR-200 family were correlated with *ACE2* expression in NSCLC^68^, SCLC and HNSCC cell lines (P<0.01; Figure 3b; Supplemental Figure 3e)^69^ and both TCGA LUAD and HNSC tumors (Figure 3b). However, forced miR-200 expression in 344SQ lung adenocarcinoma cells with high metastatic potential was not sufficient to alter *ACE2* expression (Supplemental Figure 3f2e)^70^. miR-200 family members repress *ZEB1* expression, so unsurprisingly *ACE2* is negatively correlated with *ZEB1* in cancer cell lines and tumors (Table 1). *ACE2* similarly is negatively correlated with a number of other EMT-associated genes, including *ZEB2, SNAI1, SNAI2, TWIST1*, and *CDH2* (Supplemental Table 2). While these correlations are almost invariably negative, as expected, none of these genes demonstrate a negative correlation as robust and consistent across datasets between *ACE2* as EMT score or *ZEB1*.

**Figure 3.**
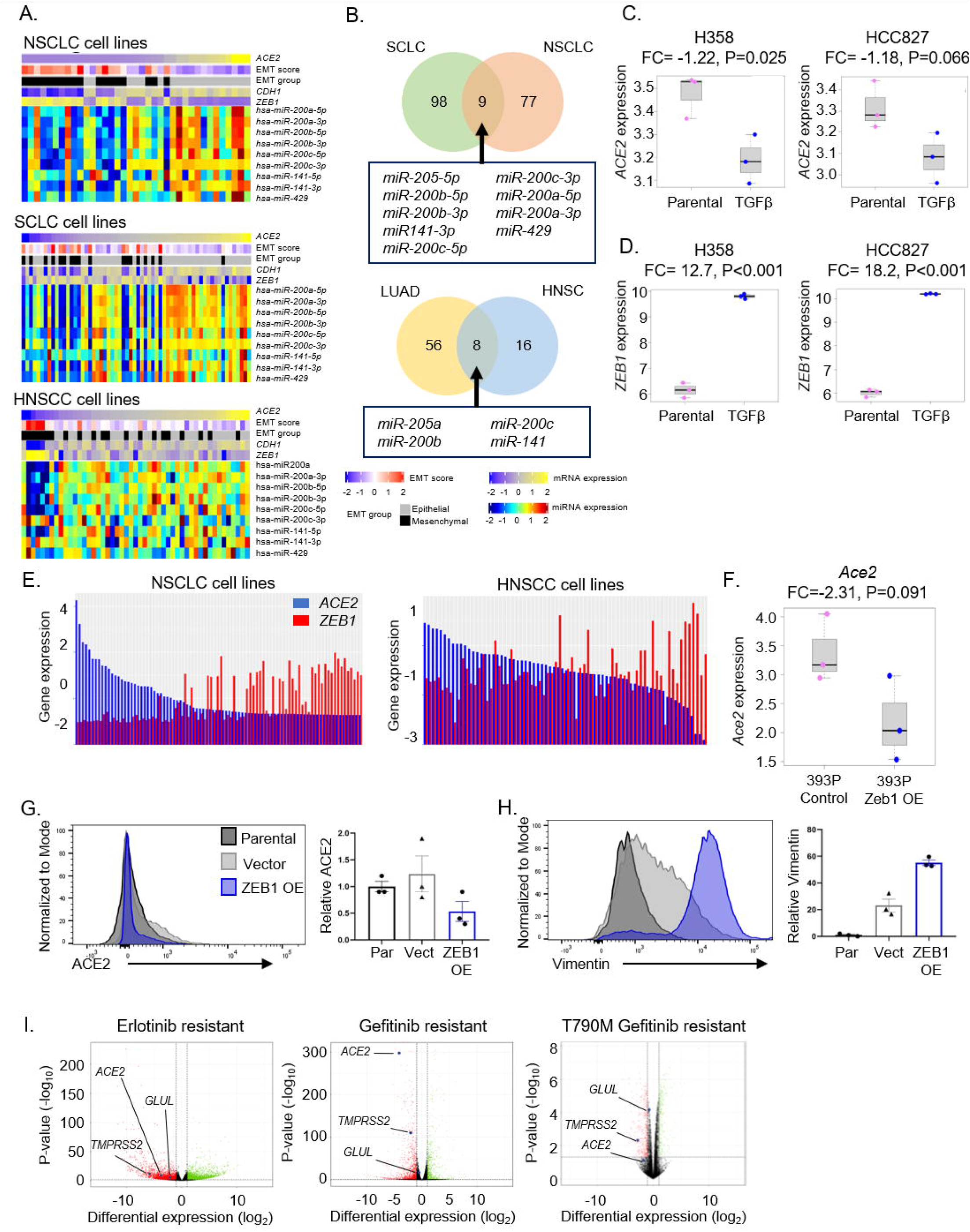
*ACE2* repression by EMT in lung cancer cells. a, *ACE2*, EMT score, EMT group (epithelial or mesenchymal), *CDH1*, and *ZEB1* expression with miR200 family miRNAs in NSCLC, SCLC and HNSCC cell lines. b, Venn diagrams demonstrating miRNAs correlated with *ACE2* expression in both NSCLC and SCLC cell lines (upper) and TCGA LUAD and HNSC tumor biopsies (lower). RNAseq data is shown as normalized relative gene expression. c,d,, Expression of *ACE2* (c) and *ZEB1*(d) following induction of EMT by TGFβ treatment in H358 and HCC827 NSCLC cell lines. e, Expression of *ACE2* and *ZEB1* in NSCLC and HNSCC cell lines. f, Expression of Ace2 in 393P murine lung adenocarcinoma cells with Zeb1 overexpression. g,h, Cell surface expression of ACE2 (g) and Vimentin (h) in HCC827 cells with constitutive overexpression of ZEB1. i, *ACE2, TMPRSS2* and *GLUL* expression in NSCLC cell lines with acquired resistance to EGFR TKIs that occurred through EMT but not when resistance occurred through EGFR T790M resistance mutations. RNAseq data is shown as normalized relative gene expression.

To determine whether transformation from an epithelial to mesenchymal phenotype (EMT) directly alters *ACE2* expression, we utilized various methods to induce EMT, including treatment with TGFβ, overexpression of ZEB1, and EGFR TKI resistance. Two highly epithelial NSCLC cell lines cultured with TGFβ for 3 to 5 weeks^71^, increased the EMT score (Supplemental Figure 4a). Induction of EMT resulted in a loss of *ACE2* in H358 and HCC827 (P=0.03 and P=0.06, respectively; Figure 3c). Additionally, both cell lines exhibited increased *ZEB1* following EMT induction (P<0.0001; Figure 3d). The experiment also included EMT induction in A549 cells, but this line is more mesenchymal with low endogenous *ACE2* expression at baseline and TGFβ was unable to reduce already minimal *ACE2* (P=0.23, data not shown).

Accordingly, NSCLC and head/neck squamous cell carcinoma (HNSCC) cell lines demonstrate an inverse correlation between *ACE2* and *ZEB1* (Figure 3e). Computational investigation of the *ACE2* promoter using the JASPAR database revealed five putative ZEB1 binding sites, suggesting that ZEB1 may directly repress *ACE2* expression^72^. Two of these predicted binding sites (ZEB1-EBOX3, ZEB1-EBOX4) were confirmed by chromatin immunoprecipitation (ChIP)seq to be functional binding sites for ZEB1 in the *ACE2* promoter in HepG2, a hepatocellular cell line (Supplemental Figure 4b). Consistent with this, induction of EMT via over-expression of Zeb1 in 393P murine lung adenocarcinoma cells results in a more than 2-fold reduction in *Ace2* expression (P=0.091; Figure 3f)^68^. Furthermore, constitutive overexpression of ZEB1 in HCC827 cells^68,73,74^ reduced cell surface expression of ACE2 compared to either the parental cell line (P=0.05) or the vector control (P=0.07; Figure 3g). As expected, overexpression of ZEB1 increased vimentin levels (compared to parental, P<0.0001 or vector control, P=0.002; Figure 3h). ACE2 is repressed by EMT and this is mediated, at least in part, by ZEB1.

*EGFR-*mutated NSCLCs are routinely and effectively treated with EGFR TKIs, but resistance inevitably develops via characteristic mechanisms including secondary *EGFR* resistance mutations (i.e. T790M) and, notably, EMT^75^. Thus, we evaluated expression of *ACE2* in *EGFR*-mutated NSCLC cells (HCC4006 and HCC827) treated with the EGFR TKI erlotinib until resistance occurred^76^. *ACE2* and *TMPRSS2* were downregulated in EGFR TKI resistant lines that had undergone EMT in comparison to parental (EGFR TKI sensitive cells) (Figure 3i).

Separately, PC-9 cells which also harbor *EGFR* activating mutations and display high baseline levels of *ACE2* were treated with the EGFR TKI gefitinib for three weeks until resistance emerged through EMT^77^ or though acquired T790M resistance mutations^78^. *ACE2* was downregulated in gefitinib-resistant cells that had undergone an EMT but not in resistant T790M+ resistant cells that maintained an epithelial phenotype. (Figure 3i). These findings are consistent with the notion of *ACE2* repression by the EMT-regulator ZEB1. In xenograft models, mice bearing PC-9 xenograft tumors treated with vehicle or the EGFR TKI osimertinib for three weeks to induce resistance through EMT were transcriptionally profiled by single-cell RNAseq^59^. *ACE2* was only detectable in vehicle-treated tumor cells, suggesting that *ACE2* expression was lost in tumors that had undergone EMT.

### Metabolic signifiers of ACE2-positive cells

In order for a virus to efficiently replicate, it must hijack the host cell’s mechanism for obtaining macromolecules including amino acids and nucleotides^79,80^. Previously, several viruses have been identified to induce host cell metabolic reprogramming to satisfy the biosynthetic and energy requirements needed for replication^81-84^. To identify metabolic features of cells able to be infected by SARS-CoV-2, metabolites associated with *ACE2* expression were investigated. *ACE2* expression correlates with glutamine in upper aerodigestive tract cell lines (P<0.01; Supplemental Figure 4d)^85^ and ACE2 levels correlate with glutamine in a subset of NSCLC cell lines (rho=0.89, P=0.02; Supplemental Figure 4e) Consistent with this finding, *ACE2* expression correlation with a list of 253 metabolism-associated genes and *GLUL*, which encodes an enzyme (glutamine synthetase) responsible for conversion of glutamate to glutamine, was identified in NSCLC, HNSCC, and SCLC cell lines (Supplemental Figure 4f) and was confirmed in the tumor data sets (Table 1). In addition to *GLUL, DERA* (deoxyribose-phosphate aldolase) and *SLC6A14* (solute carrier family 6 member 14) were identified and are implicated in nucleotide metabolism and amino acid (including glutamine) transport, respectively (Supplemental Figure 4f). Interestingly, *GLS*, which encodes the enzyme (glutaminase) that catalyzes the opposing reaction, in which glutamine is converted to glutamate, is negatively correlated with *ACE2* expression (Table 1). Next, we evaluated whether ZEB1 is a regulator of metabolism. Unlike *Ace2*, Zeb1 is unable to directly regulate *Glul* expression in murine 393P cells overexpressing Zeb1 (P=0.23, Supplemental Figure 4g). This was confirmed by *ZEB1* gene silencing and overexpression analyses (data not shown), suggesting that the metabolic changes are not directly controlled by ZEB1. Interestingly, *GLUL* was downregulated in EGFR TKI gefitinib-resistant cells that had undergone an EMT but not in resistant T790M+ resistant cells that maintained an epithelial phenotype. (Figure 3i). Similarly, *GLUL* expression was reduced following osimertinib-resistance that occurred through EMT (Supplemental Figure 4c). Taken together, glutamine synthetase-catalyzed conversion of glutamate to glutamine in epithelial cells aims to increase production of amino acids and nucleotides, while in mesenchymal cells, glutaminase converts glutamine to glutamate to preferentially shift toward the tricarboxylic acid (TCA) cycle, thus priming epithelial cells for viral infection.

### EGFR/HER family link to ACE2

NSCLC encompasses a heterogeneous group of lung cancers with unique molecular features. To determine whether any previously characterized NSCLC subsets were linked to differential *ACE2* expression, we compared common driver mutation subsets of the TCGA LUAD tumors and found that *EGFR*-mutant LUAD tumors had significantly higher *ACE2* expression (P=0.001; Supplemental Figure 4h). *EGFR*-mutated tumors have constitutive activation of the epidermal growth factor receptor (EGFR) and downstream signal transduction pathway. Accordingly, higher *ACE2* expression was associated with increased sensitivity to EGFR tyrosine kinase inhibitors (TKIs), as visualized by drug target constellation (DTECT) map (Supplemental Figure 3i). Moreover, *ACE2* expression was broadly correlated with increased expression of several EGFR/HER family members, including *EGFR, ERBB2* and *ERBB3* in cell lines and tumors (Supplemental Figure 3j), consistent with its preferential expression in epithelial cells. A similar link between EGFR inhibition and COVID-19 patients has been described previously^86^.

### Reversal of EMT

We hypothesized that bemcentinib (BGB324, or R428), an inhibitor of the mesenchymal receptor tyrosine kinase AXL with a known anti-viral effect, may offer therapeutic benefit through a reversal of EMT^87^. Bemcentinib downregulated ZEB1 expression in mesenchymal cell lines, 393P overexpressing Zeb1 (P=0.003) and Calu-1 (P=0.21) (Figure 4a). In Calu-1 cells, bemcentinib also downregulated vimentin and upregulated both E-cadherin and ACE2. Similarly, 48h of bemcentinib-treatment in HCC827 parental, vector control, and ZEB1 constitutive overexpressing lines^68,73,74^ induced ACE2, as confirmed by densitometry normalized to vinculin controls (Figure 4b). In contrast, hydroxychloroquine treatment was unable to consistently alter ACE2 or GLUL levels in a range of NSCLC cell lines (Figure 4c), indicating that this treatment is unable to alter EMT.

**Figure 4.**
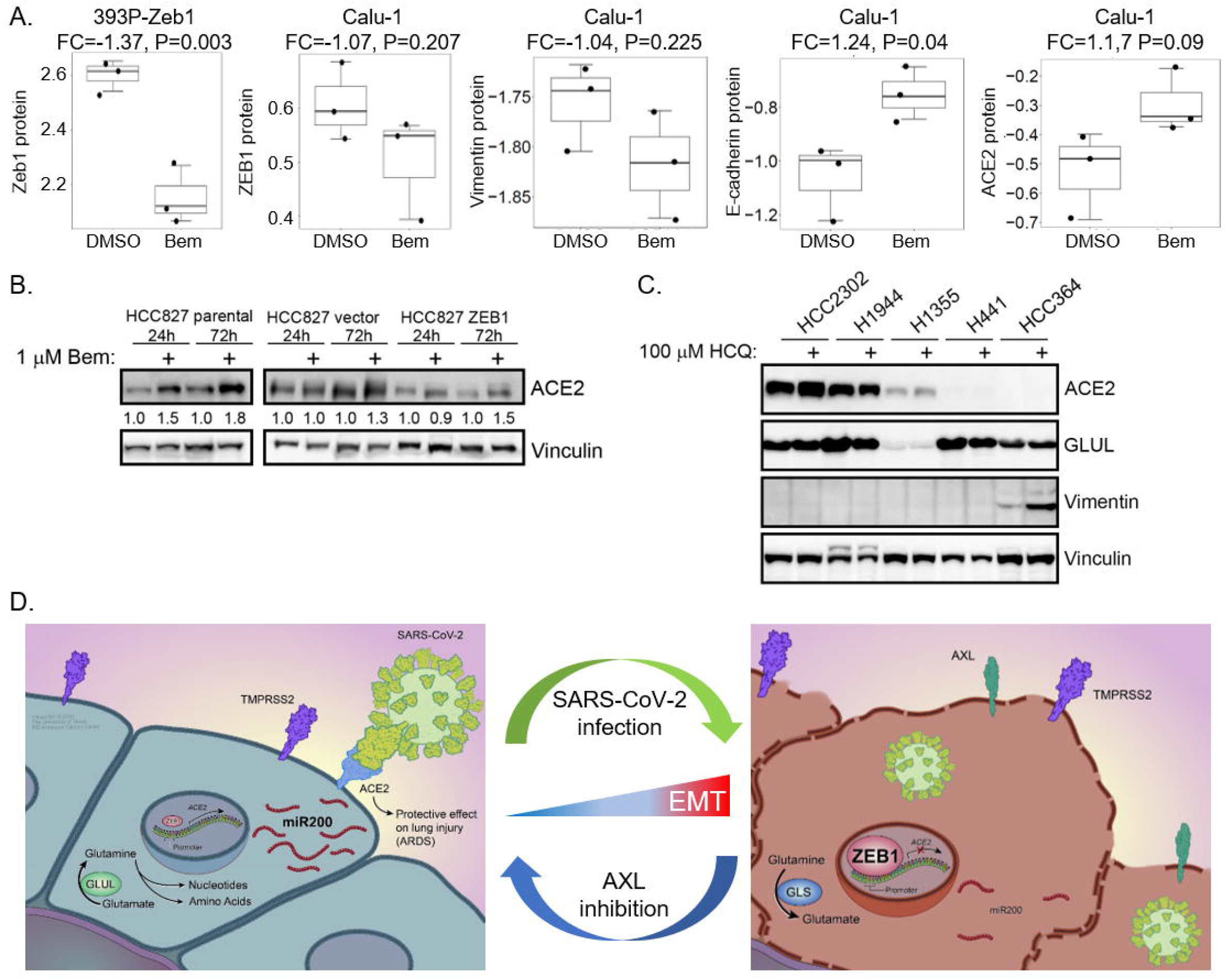
EMT modulation for treatment of SARS-CoV-2. a, ZEB1 expression following 24h of 0.5 µM bemcentinib (Bem) treatment in mesenchymal Calu-1 cells or following over-expression of Zeb1 in 393P murine cells, as well as vimentin, E-cadherin and ACE2 in Calu-1 cells. b, ACE2 levels following 24h and 72 of 1 µM Bem treatment in HCC827 cells with constitutive overexpression of ZEB1. c, ACE2, GLUL, and Vimentin levels in a subset of NSCLC cell lines following 48h treatment with 100 µM hydroxychloroquine. d, Working model demonstrating aerodigestive/respiratory cells with ACE2 expression are epithelial and metabolically primed for replication [left] while infected cells demonstrate a shift to a mesenchymal phenotype with reversal of glutamine production, a leaky cell membrane and increased ZEB1 to induce EMT. [right]. Bem = bemcentenib.

Together, these data suggest that SARS-CoV-2 infects rare, epithelial, ACE2- and TMPRSS2-positive cells in the aerodigestive and respiratory tracts that are metabolically-primed for glutamine synthesis and rapid replication (Figure 4d). Once a cell is infected with SARS-CoV-2, there is a shift to a more mesenchymal phenotype characterized by high ZEB1 and AXL, putatively lower levels of miR-200, and a decreased dependence on glutamine synthesis. Viral infection and the resultant mesenchymal shift and ZEB1 increase may have devastating implications on the risk and pathogenesis of ARDS in COVID-19 patients (Figure 4d).

## Conclusion

The ultimate public health impact of the SARS-CoV-2 pandemic has not yet been fully realized. While increased molecular understanding of the virus-host interactions is sorely needed, the impact of the pandemic itself has created practical limitations on laboratory-based research due to wide-ranging institutional restrictions. Despite these restrictions, an unprecedented collaborative research effort has ensued, with little regard for geographic and public-private boundaries that so often impose their own limitations. In the study above, we present an effort that not only incorporates our own unique cancer data sets, but also utilizes a plethora of publicly available data from basic scientists, clinicians, and bioinformaticians around the world. The linkage of regulation of *ACE2* expression and EMT, the latter an uncommon phenomenon in *normal* tissue, but well-defined in cancer cells, also highlights the value of considering malignant tissue and models for the investigation of SARS-CoV-2.

In this study, we describe a novel model for the regulation of ACE2, which others have shown is essential for the host cell entry of SARS-CoV-2^6^. In contrast to reports of patient susceptibility to SARS-CoV-2 due to age, gender, smoking history, or thoracic malignancies^4,5,88,89^ we were unable to identify consistent differences in ACE2 expression on the basis of these variables. Our data suggest that *ACE2* expression is restricted in both normal and malignant aerodigestive and respiratory tissues almost exclusively to epithelial cells, including highly specialized POU2F3-positive tuft cells, and that this restriction reflects regulatory mechanisms shared with EMT. While others have observed a putative relationship in other contexts between ACE2 and EMT, these observations were inconsistent, supporting roles for ACE2 as an inducer and an inhibitor of EMT^26,27^. In virtually all explored datasets, ranging from cancer cell lines to normal respiratory tissue, we observed a strong and consistent negative correlation between *ACE2* expression and EMT. This relationship is observed even on a cell-by-cell basis, as scRNAseq illustrates that virtually every cell with detectable *ACE2* expression is epithelial based on our published EMT score^50^.

The negative correlation between EMT and *ACE2* was not limited to expression analyses, as metabolomic analysis revealed that *ACE2* expression is strongly correlated with high levels of glutamine and the expression of *GLUL*, the enzyme responsible for the conversion of glutamate to glutamine. High levels of glutamine are characteristic of epithelial differentiation, while a shift from glutamine to glutamate is associated with EMT. We propose that SARS-CoV-2 targets these *ACE2*-expressing, metabolically-primed epithelial cells, in order to exploit the abundant nucleotides for rapid replication and viral spread. The consequence of SARS-CoV-2 infection is induction of EMT and subsequent upregulation of GLS, resulting in a depletion of glutamine following SARS-CoV-2 infection in patients^32^.

Our subsequent analyses demonstrate that, beyond mere correlations, various strategies to manipulate EMT result in predictable alterations in the expression of *ACE2*. For example, treatment of cells with TGFβ or overexpression of ZEB1 induces EMT *and* decreased expression of *ACE2*. Similarly, in *EGFR*-mutant lung adenocarcinoma, in which there is relatively higher *ACE2* expression, prolonged treatment with an *EGFR*-targeting TKI may result in multiple mechanisms of therapeutic resistance, of which EMT is a common one. Our results demonstrate that *ACE2* is downregulated when EMT is the underlying resistance mechanism, but not when resistance emerges due to development of secondary resistance mutations in *EGFR*. Coincident with the loss of *ACE2* expression is the downregulation of *GLUL*, to promote glutamate and enable production of TCA intermediates and energy.

EMT is a complex process regulated by an intricate molecular network and we hoped to discern whether any of the specific pathway(s) that regulate EMT also mediate *ACE2* expression. An investigation into one of the transcriptional mechanisms underlying EMT and its association with *ACE2* revealed consistent positive correlations across tissue and sample types with the miR-200 family of microRNAs. The miR-200 family has a well-established role as an inhibitor of EMT via the downregulation of the transcriptional repressor ZEB1. Predictably, a strong negative correlation was observed between *ACE2* and *ZEB1*. The repression of ACE2 by ZEB1 is mediated via two repressor binding sites for within the *ACE2* promoter and forced over-expression of *ZEB1* results in a more than two-fold decrease in *ACE2* expression, but not *GLUL*. ZEB1 regulation occurs via SNAI1 and/or TWIST1 while SNAI1 induces nuclear translocation of ETS1, which is required for ZEB1 expression^90^. It is likely that EMT induction of these genes by SARS-CoV-2 results in increased ZEB1. Finally, we observed that SARS-CoV-2 infected lung cancer cell lines, bronchial organoids and patient nasopharangeal swabs demonstrate significant shifts toward several EMT-like features including upregulation of *ZEB1* and downregulation of *EPCAM*.

Together, our data, in the context of the ever-growing literature on SARS-CoV-2, suggest that *ACE2* expression is necessary for initial viral entry and, therefore, is relatively high in those cells infected by the virus. These cells are epithelial and, in our data sets, unexpectedly rare, considering the devastating impact of this infection. Following viral entry, however, SARS-CoV-2 infection induces molecular changes within the cells that are reminiscent of EMT – especially the increased expression of *ZEB1*. These SARS-CoV-2-induced changes, compounded by the downregulation of genes, including tight junction components, which play a critical role in restricting epithelial and endothelial permeability, exposes respiratory cells to increased risk of edema and ARDS. Interestingly, previous reports with SARS-CoV(−1) demonstrate that lung cells, including alveolar epithelial cells, express higher levels of TGFβ, a key regulator of EMT^91^. It is not surprising that a similar mechanism would occur with a similar coronavirus, such as SARS-CoV-2.

While these data demonstrate an intriguing mechanism for the molecular virus-host interaction, it is critical to consider how these observations may be harnessed to develop strategies to improve patient outcomes. This, as always, is complex. Would a therapeutic effort focused on prevention of the initial infection by decreasing ACE2 levels, for example with an inhibitor of the miR-200 family, have unforeseen consequences such as increased risk of ARDS? Similarly, would therapeutic efforts post-infection aimed at preventing SARS-CoV-2 induced ACE2 downregulation, for example with a miR-200 mimetic, serve to place adjacent, newly ACE2-positive cells at risk for infection? Several large cohort observational studies on ACE inhibitors and angiotensin II receptor blockers (ARBs), which upregulate ACE2 expression, did not show association with either SARS-Cov-2 infection or risk of severe COVID-19^92^. Interestingly, a human recombinant soluble ACE2 which has already been tested in phase 1 and phase 2 clinical trials for ARDS showed inhibition of SARS-CoV-2 infection in human capillary organoids and human kidney organoids^93^.

Based on the findings in our study, an attractive alternative therapeutic strategy would be to reverse EMT with an AXL inhibitor. AXL is overexpressed in several solid and hematologic malignancies, including non-small cell lung cancer (NSCLC), acute myeloid leukemia (AML), breast and prostate cancer, among others. In the context of malignancy, AXL overexpression can drive EMT, tumor angiogenesis, and decreased antitumor immune response^61^. Furthermore, AXL inhibitor, gilteritinib, was identified to have anti-viral activity against SARS-CoV-2^94^. Bemcentinib, a highly selective and potent inhibitor of AXL, currently being tested in phase 2 trials in various malignancies including NSCLC, AML and breast cancer. In preclinical models, bemcentinib has already demonstrated potent anti-viral activity in Ebola and Zika viruses^14,95^. Recent data suggests that bemcentinib inhibits SARS-CoV-2 viral entry into cells and also prevents inhibition of type I interferon, which is the host cells’ innate antiviral immune response. In contrast to AXL inhibition, hydroxychloroquine, an alternative treatment for COVID-19 patients, does not demonstrate any ability to reverse EMT. Accordingly, bemcentinib was recently fast-tracked as the first potential treatment for assessment in the United Kingdom’s <u>AC</u>celerating <u>CO</u>VID-19 <u>R</u>esearch & <u>D</u>evelopment (ACCORD) multicenter, randomized Phase II trial^17^. However, the trial has ceased in the UK due to low numbers of COVID-19 patients and the company is seeking regulatory approval to conduct a similar trial in a country with high COVID-19 incidence. Nevertheless, these data reveal EMT, along with associated proteins such as ZEB1 and AXL, as novel therapeutic targets of interest for combating COVID-19.

## Methods

### Transcriptional sequencing datasets

RNAseq analysis datasets from normal tissues were obtained from GTEx Portal (http://www.gtexportal.org) and tumor samples were retrieved from published and unpublished datasets in multiple human tissues, including TCGA LUAD^43^, TCGA HNSC^44^, TCGA LUSC, PROSPECT^45^, BATTLE 1^46^, and BATTLE 2^47^. Cell line RNAseq data were obtained from multiple sources, including CCLE upper aerodigestive tract (n=32), LCNEC cell lines (n=15)^42^, as well as NSCLC (n=88)^37,38^, HNSCC (n=70)^40^, and SCLC (n=62)^48^.

Transcriptional data from experimental datasets were publicly available, including EMT induction by three weeks of TGFβ^71^, 393P murine cells with Zeb1 overexpression^68^, erlotinib-resistance via EMT in *EGFR*-mutated HCC4006 and HCC827 NSCLC cell lines^76^, gefitinib-resistant PC-9 cells via EMT^77^, T790M-mediated gefitinib-resistance in PC-9 cells^78^, Calu-3 or A549 transduced with a vector expressing human ACE2 were mock infected or infected with SARS-CoV-2 (USA-WA1/2020)^60^, forced miR-200 expression in 344SQ lung adenocarcinoma cells with high metastatic potential^70^, human bronchial organoids were infected with SARS-CoV-2 for five days^65^, and nasopharangeal swabs from patients with positive or negative SARS-CoV-2 PCR results^66^.

Single-cell RNAseq data were obtained from publicly available sources, including bronchial epithelial cells obtained by bronchial brushings from six never smokers and six current smokers^96^, oral cavity tumors from 18 tumors^55^, five normal donor and eight fibrotic lungs^57^, freshly resected parenchymal lung tissue from three patients^58^, bronchial biopsy, nasal brushing, nasal turbinate specimens (GSE121600), and PC-9 *EGFR*-mutant NSCLC cell xenograft tumors treated with vehicle or osimertinib for three weeks^59^.

ChIPseq analysis of ZEB1 binding in the *ACE2* promoter of HepG2, human hepatocyte carcinoma cells, as previously described^97^.

### Flow Cytometry

One million cells each for HCC2302, H3255, HCC827, H1944, H441, H2023, HCC364, A549, H1355, HCC827 vector, and HCC827 ZEB1 were surfaced stained with ACE2 (Santa Cruz; sc-390851) then fixed in 2% PFA and permeabilized using 1X Intracellular Staining Perm Wash Buffer (BioLegend; 421002) prior to intracellular staining with Vimentin (BD Pharmingen; 562338). Samples were analyzed on a BD LSRFortessa Flow Cytometer and data was analyzed using FlowJo 10.7.1. Graphs and statistical analysis were done using GraphPad Prism 8.

### Metabolite Assay

One million cells each for 393P ZEB1 (DMSO 16h), 393P ZEB1 (DOX 16h), H441, H1944, HCC2302, HCC827, H2023, and H1355 were harvested. Cell lysates were prepared by addition of 0.3N hydrochloric acid then 450nM Tris (pH=8) and metabolites were analyzed using the Glutamine/Glutamate Glo Assay (Promega; J8021) per manufacturer’s instruction.

### Drug treatment of cells

393P-Zeb1 inducible murine NSCLC cells were treated with AXL inhibitor, bemcentinib (0.5 uM), or DMSO for total 24h (8h pretreatment with bemcentinib or DMSO, followed by induction of Zeb1 with doxycycline (2 ug/mL) and an additional 16h drug treatment). Five NSCLC cell lines with varying levels of ACE2 were treated with hydroxychloroquine sulfate (Sigma) for 48h. Cell lysates were harvested for western blot or RPPA analysis. Antibodies used for western analysis include ACE2 (MA5-32307, ThermoFisher), GLUL (80636, Cell Signaling Technology), Vimentin (3932, Cell Signaling Technology), ZEB1 (3396, Cell Signaling Technology), and vinculin (Sigma, V9131) as a loading control.

### RPPA

Protein lysates from TCGA LUAD, NSCLC cell lines, and Calu-1 or 393P cells treated with bemcentinib were quantified and protein arrays were printed and stained^98^. Images were quantified with MicroVigene 4.0 (VigeneTech, Carlisle, MA). The spot-level raw data were processed with the R package SuperCurve suite, which returns the estimated protein concentration (raw concentration) and a quality control score for each slide, as described previously^98^. Only slides with a quality control score of >0.8 were used for downstream analysis. The raw concentration data were normalized by median-centering each sample across all the proteins to correct loading bias. *microRNA Arrays* Total RNA from 55 NSCLC, 30 SCLC, and 49 HNSCC cell lines were analyzed with Affymetrix miRNA 4.0 arrays. The expression data of a curated list of miR-200 family members (hsa-miR-200b-5p, hsa-miR-200b-3p, hsa-miR-200c-5p, hsa-miR-200c-3p, hsa-miR-200a-5p, hsa-miR-200a-3p, hsa-miR-429, hsa-miR-141-5p, hsa-miR-141-3p) known to be involved with EMT in NSCLC^67^ were compared with ACE2 expression data.

### Metabolite analyses

A total of 225 metabolites were profiled in CCLE upper aerodigestive tract cell lines using liquid chromatography–mass spectrometry (LC-MS)^85^. We compared *ACE2* mRNA expression to abundance of metabolites.

### Computational prediction of binding sites

To generate the predicted ZEB1 (E-Box) binding sites on the ACE2 promoter, the promoter sequence for human ACE2 was downloaded and used in the matrix profile search on the JASPAR web portal^72^. For the search, the vertebrate database selecting ZEB1 binding motifs were selected to search through the promoter sequence of ACE2 with a relative profile score threshold score cutoff of 80% or higher. The resulting sites were ranked and accordingly the highest-scoring motifs were annotated on the promoter segment using Snapgene.

### Single-cell RNAseq Analysis

Raw data for single cell datasets were downloaded from GEO data base and processed using the Seurat Package v2.3.1^99^. First the raw read counts were normalized and scaled using “NormalizeData” and “ScaleData” function. The most variable genes were selected using the “FindVariableGenes” function with default parameters, and principle component analysis (PCA) were performed on these variable genes. We selected the first N principle components that account for at least 80% of the total variances to perform the tSNE transformation. For each genes, the expression status is defined as positive if the cell has non-zero expression value, or negative otherwise. For GSE122960, samples were grouped in to donor samples (including GSM3489182, GSM3489187, GSM3489189, GSM3489191 and GSM3489193) and fibrotic samples (including GSM3489183, GSM3489184, GSM3489188, GSM3489190, GSM3489192, GSM3489194, GSM3489196 and GSM3489198). Each group was subsampled to 20,000 total cells. EMT score was calculated based on the EMT signature as described previously^49,100^. Cell-specific expression of *SFTPC* (alveolar type II cells), *FOXJI, PIFO* (ciliated cells), and *SCGB1A1* (club cells) were visualized in tSNE feature plots to identify epithelial clusters containing *ACE2*-positive cells.

### Statistical Analyses

All statistic and bioinformatics analyses were performed using R. Paired and un-paired two-sample t-tests were used for two group comparisons on paired and unpaired experimental design experiments. Pearson and Spearman correlations were used for correlating genomic and proteomic measurements, as well as correlating drug-screening data. In all the analyses, p<0.05 was considered statistically significant.

The drug-target constellation map (DTECT map) was generated based on drug screening data using a suite of R packages, including ggplot2, ggraph and igraph. A list of drugs was selected based on Spearman coefficient (>0.3) and p value (<0.1) and only targets with more than 5 associated drugs were shown in the figure.

### Data Set Availability

Publicly available data were obtained from GEO datasets: GSE122960, GSE131391, GSE103322, GSE42127, GSE122960, GSE130148, GSE138693, GSE121600, GSE49644, GSE61395, GSE114647, GSE75602, GSE147507, GSE15741, GSE150819, GSE154770, GSE32465 (GSM1010809).

## Acknowledgements

We wish to acknowledge the patients impacted by the COVID-19 pandemic, along with their loved ones, as well as the first responders, healthcare staff, and essential workers who worked tirelessly and selflessly during this devastating period. We thank C.E.S. for graphical elements and David M. Aten for assistance with medical illustrations. We would like to thank P.M.G., M.P.B, W.A.B, G.D.S, P.W.S., R.A.S., R.R.S., R.A.D., B.A.D., V.J.M.C., S.C., S.C., J.P., S.I.1., S.I.2., and C.J.C. for intellectual support provided during quarantine. The work was supported by NIH/NCI R50-CA243698 (C.A.S.), NIH/NCI R01-CA207295 (L.A.B.), NIH/NCI U01-CA213273 (L.A.B., J.V.H.), CCSG P30-CA01667 (L.A.B.), University of Texas SPORE in Lung Cancer P5-CA070907 (L.A.B., D.L.G., J.V.H., C.M.G., J.W., K.R., K.R.C.), the Department of Defense (LC170171; L.A.B.), Khalifa Bin Zayed Al Nahyan Foundation (C.M.G.), CPRIT Research Training Program RP170067 (E.M.P.), The IASLC (K.R.C.), The University of Texas MD Anderson Cancer Center-Oropharynx Cancer Program generously supported by Mr. and Mrs. Charles W. Stiefel (F.M.J.), through generous philanthropic contributions to The University of Texas MD Anderson Lung Cancer Moon Shot Program, The Jane Ford Petrin Fund, The Andrew Sabin Family Fellowship, and The Rexanna Foundation for Fighting Lung Cancer.

## Author Contributions

C.A.S., C.M.G., and L.A.B. conceived the project, analyzed and interpreted the data, and wrote the manuscript; K.R. and K.R.C. performed experiments and interpreted results; R.J.C., M.N., E.M.P, C.M.D.C., K.K., interpreted results; S.H., S.K., L.D., Q.W., L.S., Y.X., and J.W. contributed to the analysis and interpretation of data; C.P., F.M.J., J.Z., H.K., J.M., D.L.G, and J.V.H. contributed to the acquisition of data, administrative, and/or material support. All authors contributed to the writing, review, and/or revision of the manuscript.

## Supplemental Figures

Supplemental Figure 1. ACE2 expression in select populations and cell-types. a, *ACE2* mRNA expression in a panel of non-malignant tissue types, including lung and salivary gland. b, Expression of *ACE2* and EMT score in TCGA LUAD biopsies and normal adjacent tissue. c, Forest plot demonstrating no difference in *ACE2* expression based on smoking status, gender or age. d, Vimentin expression by flow cytometry in a subset of NSCLC cell lines.

Supplemental Figure 2. *ACE2* expression is regulated by mechanisms similar to those governing EMT, which are co-opted by SARS-CoV-2 infection. a, EMT score expression for each *ACE2*-positive cell present in pooled donor or fibrotic lungs. b, EMT score values for each *ACE2*-positive cell present in oral cavity tumors indicated by color. c, tSNE visualization of epithelial cells from normal and fibrotic lungs demonstrating expression of cell type-specific genes, as well as *ACE2*. RNAseq data is shown as normalized relative gene expression.

Supplemental Figure 3. SARS-CoV-2 induced changes. a, Effects of 24h SARS-CoV-2 infection of A549 cells on *ZEB1, AXL, EPCAM* and *CLDN2* expression in A549 cells with undetectable ACE2 levels at the time of infection. b, Effects of SARS-CoV-2 infection of Calu-3 or A549+ACE2 for 24h on *GLUL* expression. c, Effects of SARS-CoV-2 infection in patient nasal swabs for *EPCAM, ZEB1, AXL* and EMT score broken down by viral load. d, Effects of SARS-CoV-2 infection on *GLS* expression. *P<0.10, **P<0.01,***P<0.005, ****P<0.0001. e, *ACE2* mRNA correlation with miRNAs in SCLC and NSCLC cell lines. Blue boxes surround the miR200 family members. f, *ACE2* expression in forced miR-200 expression in 344SQ lung adenocarcinoma cells. RNAseq data is shown as normalized relative gene expression.

Supplemental Figure 4. Regulation of ACE2 by EMT. a, EMT score following EMT induction by TGFβ. b, ZEB1 binding sites present in the promoter of *ACE2* in HepG2 cells by ChIPseq. c, *ACE2*- and *GLUL*-positive cells are reduced in PC-9 cells with EGFR TKI resistance via EMT. d, *ACE2* expression is correlated with specific metabolites in aerodigestive tract cells. e, ACE2 protein is correlated with glutamine in NSCLC cell lines. f, Venn diagrams demonstrating metabolic genes correlated with *ACE2* in NSCLC, HNSCC, and SCLC cell lines. g, *Glul* expression following overexpression of Zeb1 in 393P murine lung adenocarcinoma cells. h, *ACE2* expression is higher in TCGA LUAD tumor biopsies with *EGFR* mutations. i, *ACE2* expression is correlated with response to a large number of EGFR TKIs, including those that are FDA approved (underlined). j, High *ACE2* is associated with high expression of EGFR/HER family members, including *EGFR, ERBB2, ERBB3*, in a number of aerodigestive and respiratory tract cell lines.

Supplemental Table 1. *ACE2* co-expression with *TMPRSS2* and *POU2F3* in normal and fibrotic lungs. *P<0.05, **P<0.01, ***P<0.001.

Supplementary Table 2. ACE2 rho correlation values in aerodigestive and respiratory tract cell line and tumor biopsy specimens. *P<0.05, **P<0.01, ***P<0.001.

## References

1. Wang, C., Horby, P.W., Hayden, F.G. & Gao, G.F. A novel coronavirus outbreak of global health concern. Lancet 395, 470–473 (2020).

2. Zhu, N., et al. A Novel Coronavirus from Patients with Pneumonia in China, 2019. N Engl J Med 382, 727–733 (2020).

3. Zhou, P., et al. A pneumonia outbreak associated with a new coronavirus of probable bat origin. Nature 579, 270–273 (2020).

4. Dai, M., et al. Patients with Cancer Appear More Vulnerable to SARS-COV-2: A Multicenter Study during the COVID-19 Outbreak. Cancer Discov (2020).

5. Yu, J., Ouyang, W., Chua, M.L.K. & Xie, C. SARS-CoV-2 Transmission in Patients With Cancer at a Tertiary Care Hospital in Wuhan, China. JAMA Oncol (2020).

6. Hoffmann, M., et al. SARS-CoV-2 Cell Entry Depends on ACE2 and TMPRSS2 and Is Blocked by a Clinically Proven Protease Inhibitor. Cell 181, 271–280 e278 (2020).

7. Cao, Y., et al. Comparative genetic analysis of the novel coronavirus (2019-nCoV/SARS-CoV-2) receptor ACE2 in different populations. Cell Discov 6, 11 (2020).

8. Imai, Y., et al. Angiotensin-converting enzyme 2 protects from severe acute lung failure. Nature 436, 112–116 (2005).

9. Yang, X., et al. Clinical course and outcomes of critically ill patients with SARS-CoV-2 pneumonia in Wuhan, China: a single-centered, retrospective, observational study. Lancet Respir Med 8, 475–481 (2020).

10. Kuba, K., et al. A crucial role of angiotensin converting enzyme 2 (ACE2) in SARS coronavirus-induced lung injury. Nat Med 11, 875–879 (2005).

11. Matsuyama, S., et al. Enhanced isolation of SARS-CoV-2 by TMPRSS2-expressing cells. Proc Natl Acad Sci U S A 117, 7001–7003 (2020).

12. Ziegler, C.G.K., et al. SARS-CoV-2 Receptor ACE2 Is an Interferon-Stimulated Gene in Human Airway Epithelial Cells and Is Detected in Specific Cell Subsets across Tissues. Cell 181, 1016–1035 e1019 (2020).

13. Brindley, M.A., et al. Tyrosine kinase receptor Axl enhances entry of Zaire ebolavirus without direct interactions with the viral glycoprotein. Virology 415, 83–94 (2011).

14. Meertens, L., et al. Axl Mediates ZIKA Virus Entry in Human Glial Cells and Modulates Innate Immune Responses. Cell Rep 18, 324–333 (2017).

15. Chen, J., et al. AXL promotes Zika virus infection in astrocytes by antagonizing type I interferon signalling. Nat Microbiol 3, 302–309 (2018).

16. University of Iowa virology research helps facilitate new clinical trial for COVID-19. (ed. Iowa, U.o.) (https://uihc.org/news/university-iowa-virology-research-helps-facilitate-new-clinical-trial-covid-19, 2020).

17. BerGenBio. BERGENBIO’S BEMCENTINIB SELECTED TO BE FAST-TRACKED AS POTENTIAL TREATMENT FOR COVID-19 THROUGH NEW NATIONAL UK GOVERNMENT CLINICAL TRIAL INITIATIVE. (https://www.bergenbio.com/bergenbios-bemcentinib-selected-to-be-fast-tracked-as-potential-treatment-for-covid-19-through-new-national-uk-government-clinical-trial-initiative/, 2020).

18. Spinato, G., et al. Alterations in Smell or Taste in Mildly Symptomatic Outpatients With SARS-CoV-2 Infection. JAMA (2020).

19. Xydakis, M.S., et al. Smell and taste dysfunction in patients with COVID-19. Lancet Infect Dis (2020).

20. Howitt, M.R., et al. Tuft cells, taste-chemosensory cells, orchestrate parasite type 2 immunity in the gut. Science 351, 1329–1333 (2016).

21. O’Leary, C.E., Schneider, C. & Locksley, R.M. Tuft Cells-Systemically Dispersed Sensory Epithelia Integrating Immune and Neural Circuitry. Annu Rev Immunol 37, 47–72 (2019).

22. Matsumoto, I., Ohmoto, M., Narukawa, M., Yoshihara, Y. & Abe, K. Skn-1a (Pou2f3) specifies taste receptor cell lineage. Nat Neurosci 14, 685–687 (2011).

23. Ohmoto, M., et al. Pou2f3/Skn-1a is necessary for the generation or differentiation of solitary chemosensory cells in the anterior nasal cavity. Biosci Biotechnol Biochem 77, 2154–2156 (2013).

24. Yamashita, J., Ohmoto, M., Yamaguchi, T., Matsumoto, I. & Hirota, J. Skn-1a/Pou2f3 functions as a master regulator to generate Trpm5-expressing chemosensory cells in mice. PLoS One 12, e0189340 (2017).

25. Huang, Y.H., et al. POU2F3 is a master regulator of a tuft cell-like variant of small cell lung cancer. Genes Dev 32, 915–928 (2018).

26. Burns, W.C., Velkoska, E., Dean, R., Burrell, L.M. & Thomas, M.C. Angiotensin II mediates epithelial-to-mesenchymal transformation in tubular cells by ANG 1-7/MAS-1-dependent pathways. Am J Physiol Renal Physiol 299, F585–593 (2010).

27. Qian, Y.R., et al. Angiotensin-converting enzyme 2 attenuates the metastasis of non-small cell lung cancer through inhibition of epithelial-mesenchymal transition. Oncol Rep 29, 2408–2414 (2013).

28. Dongre, A. & Weinberg, R.A. New insights into the mechanisms of epithelial-mesenchymal transition and implications for cancer. Nat Rev Mol Cell Biol 20, 69–84 (2019).

29. Park, S.M., Gaur, A.B., Lengyel, E. & Peter, M.E. The miR-200 family determines the epithelial phenotype of cancer cells by targeting the E-cadherin repressors ZEB1 and ZEB2. Genes Dev 22, 894–907 (2008).

30. Title, A.C., et al. Genetic dissection of the miR-200-Zeb1 axis reveals its importance in tumor differentiation and invasion. Nat Commun 9, 4671 (2018).

31. Ramirez-Pena, E., et al. The Epithelial to Mesenchymal Transition Promotes Glutamine Independence by Suppressing GLS2 Expression. Cancers (Basel) 11(2019).

32. Shen, B., et al. Proteomic and Metabolomic Characterization of COVID-19 Patient Sera. Cell 182, 59–72 e15 (2020).

33. Kim, K.K., et al. Alveolar epithelial cell mesenchymal transition develops in vivo during pulmonary fibrosis and is regulated by the extracellular matrix. Proc Natl Acad Sci U S A 103, 13180–13185 (2006).

34. Gouda, M.M., Shaikh, S.B. & Bhandary, Y.P. Inflammatory and Fibrinolytic System in Acute Respiratory Distress Syndrome. Lung 196, 609–616 (2018).

35. Cao, Y., et al. miR-200b/c attenuates lipopolysaccharide-induced early pulmonary fibrosis by targeting ZEB1/2 via p38 MAPK and TGF-beta/smad3 signaling pathways. Lab Invest 98, 339–359 (2018).

36. Huang, C., et al. Clinical features of patients infected with 2019 novel coronavirus in Wuhan, China. Lancet 395, 497–506 (2020).

37. Zhou, B.B., et al. Targeting ADAM-mediated ligand cleavage to inhibit HER3 and EGFR pathways in non-small cell lung cancer. Cancer Cell 10, 39–50 (2006).

38. Lockwood, W.W., et al. DNA amplification is a ubiquitous mechanism of oncogene activation in lung and other cancers. Oncogene 27, 4615–4624 (2008).

39. Kalu, N.N., et al. Genomic characterization of human papillomavirus-positive and - negative human squamous cell cancer cell lines. Oncotarget 8, 86369–86383 (2017).

40. Kalu, N.N., et al. Comprehensive pharmacogenomic profiling of human papillomavirus-positive and -negative squamous cell carcinoma identifies sensitivity to aurora kinase inhibition in KMT2D mutants. Cancer Lett 431, 64–72 (2018).

41. Barretina, J., et al. The Cancer Cell Line Encyclopedia enables predictive modelling of anticancer drug sensitivity. Nature 483, 603–607 (2012).

42. Barretina, J., et al. Addendum: The Cancer Cell Line Encyclopedia enables predictive modelling of anticancer drug sensitivity. Nature 565, E5–E6 (2019).

43. Cancer Genome Atlas Research, N. Comprehensive molecular profiling of lung adenocarcinoma. Nature 511, 543–550 (2014).

44. Cancer Genome Atlas, N. Comprehensive genomic characterization of head and neck squamous cell carcinomas. Nature 517, 576–582 (2015).

45. Tang, H., et al. A 12-gene set predicts survival benefits from adjuvant chemotherapy in non-small cell lung cancer patients. Clin Cancer Res 19, 1577–1586 (2013).

46. Kim, E.S., et al. The BATTLE trial: personalizing therapy for lung cancer. Cancer Discov 1, 44–53 (2011).

47. Papadimitrakopoulou, V., et al. The BATTLE-2 Study: A Biomarker-Integrated Targeted Therapy Study in Previously Treated Patients With Advanced Non-Small-Cell Lung Cancer. J Clin Oncol 34, 3638–3647 (2016).

48. Gay, C.M., et al. Patterns of transcription factor programs and immune pathway activation define four major subtypes of SCLC with distinct therapeutic vulnerabilities. Cancer Cell in press(2020).

49. Byers, L.A., et al. An epithelial-mesenchymal transition gene signature predicts resistance to EGFR and PI3K inhibitors and identifies Axl as a therapeutic target for overcoming EGFR inhibitor resistance. Clin Cancer Res 19, 279–290 (2013).

50. Mak, M.P., et al. A Patient-Derived, Pan-Cancer EMT Signature Identifies Global Molecular Alterations and Immune Target Enrichment Following Epithelial-to-Mesenchymal Transition. Clin Cancer Res 22, 609–620 (2016).

51. Hmeljak, J., et al. Integrative Molecular Characterization of Malignant Pleural Mesothelioma. Cancer Discov 8, 1548–1565 (2018).

52. Alqahtani, J.S., et al. Prevalence, Severity and Mortality associated with COPD and Smoking in patients with COVID-19: A Rapid Systematic Review and Meta-Analysis. PLoS One 15, e0233147 (2020).

53. Di Stadio, A., Ricci, G., Greco, A., de Vincentiis, M. & Ralli, M. Mortality rate and gender differences in COVID-19 patients dying in Italy: A comparison with other countries. Eur Rev Med Pharmacol Sci 24, 4066–4067 (2020).

54. Onder, G., Rezza, G. & Brusaferro, S. Case-Fatality Rate and Characteristics of Patients Dying in Relation to COVID-19 in Italy. JAMA (2020).

55. Puram, S.V., et al. Single-Cell Transcriptomic Analysis of Primary and Metastatic Tumor Ecosystems in Head and Neck Cancer. Cell 171, 1611–1624 e1624 (2017).

56. Yaegashi, H. & Takahashi, T. [Medical changes in preformed host arteries involved in a tumor--a morphometric study of gastric, colonic, and pancreatic carcinomas in man]. Gan No Rinsho 34, 1555–1560 (1988).

57. Reyfman, P.A., et al. Single-Cell Transcriptomic Analysis of Human Lung Provides Insights into the Pathobiology of Pulmonary Fibrosis. Am J Respir Crit Care Med 199, 1517–1536 (2019).

58. Vieira Braga, F.A., et al. A cellular census of human lungs identifies novel cell states in health and in asthma. Nat Med 25, 1153–1163 (2019).

59. Kurppa, K.J., et al. Treatment-Induced Tumor Dormancy through YAP-Mediated Transcriptional Reprogramming of the Apoptotic Pathway. Cancer Cell 37, 104–122 e112 (2020).

60. Blanco-Melo, D., et al. Imbalanced Host Response to SARS-CoV-2 Drives Development of COVID-19. Cell (2020).

61. Gay, C.M., Balaji, K. & Byers, L.A. Giving AXL the axe: targeting AXL in human malignancy. Br J Cancer 116, 415–423 (2017).

62. Ikenouchi, J., Matsuda, M., Furuse, M. & Tsukita, S. Regulation of tight junctions during the epithelium-mesenchyme transition: direct repression of the gene expression of claudins/occludin by Snail. J Cell Sci 116, 1959–1967 (2003).

63. Herrero, R., Sanchez, G. & Lorente, J.A. New insights into the mechanisms of pulmonary edema in acute lung injury. Ann Transl Med 6, 32 (2018).

64. Overgaard, C.E., Mitchell, L.A. & Koval, M. Roles for claudins in alveolar epithelial barrier function. Ann N Y Acad Sci 1257, 167–174 (2012).

65. Suzuki, T., et al. Generation of human bronchial organoids for SARS-CoV-2 research. bioRxiv (2020).

66. Lieberman, N.A.P., et al. In vivo antiviral host transcriptional response to SARS-CoV-2 by viral load, sex, and age. PLoS Biol 18, e3000849 (2020).

67. Chen, L., et al. Metastasis is regulated via microRNA-200/ZEB1 axis control of tumour cell PD-L1 expression and intratumoral immunosuppression. Nat Commun 5, 5241 (2014).

68. Kundu, S.T., et al. The miR-200 family and the miR-183∼96∼182 cluster target Foxf2 to inhibit invasion and metastasis in lung cancers. Oncogene 35, 173–186 (2016).

69. Allison Stewart, C., et al. Dynamic variations in epithelial-to-mesenchymal transition (EMT), ATM, and SLFN11 govern response to PARP inhibitors and cisplatin in small cell lung cancer. Oncotarget 8, 28575–28587 (2017).

70. Gibbons, D.L., et al. Contextual extracellular cues promote tumor cell EMT and metastasis by regulating miR-200 family expression. Genes Dev 23, 2140–2151 (2009).

71. Sun, Y., et al. Metabolic and transcriptional profiling reveals pyruvate dehydrogenase kinase 4 as a mediator of epithelial-mesenchymal transition and drug resistance in tumor cells. Cancer Metab 2, 20 (2014).

72. Fornes, O., et al. JASPAR 2020: update of the open-access database of transcription factor binding profiles. Nucleic Acids Res 48, D87–D92 (2020).

73. Peng, D.H., et al. ZEB1 suppression sensitizes KRAS mutant cancers to MEK inhibition by an IL17RD-dependent mechanism. Sci Transl Med 11(2019).

74. Peng, D.H., et al. ZEB1 induces LOXL2-mediated collagen stabilization and deposition in the extracellular matrix to drive lung cancer invasion and metastasis. Oncogene 36, 1925–1938 (2017).

75. Le, X., et al. Landscape of EGFR-Dependent and -Independent Resistance Mechanisms to Osimertinib and Continuation Therapy Beyond Progression in EGFR-Mutant NSCLC. Clin Cancer Res 24, 6195–6203 (2018).

76. Nilsson, M.B., et al. The YAP/FOXM1 axis regulates EMT-associated EGFR inhibitor resistance and increased expression of spindle assembly checkpoint components. Sci Transl Med ((in review)).

77. Raoof, S., et al. Targeting FGFR overcomes EMT-mediated resistance in EGFR mutant non-small cell lung cancer. Oncogene 38, 6399–6413 (2019).

78. Hata, A.N., et al. Tumor cells can follow distinct evolutionary paths to become resistant to epidermal growth factor receptor inhibition. Nat Med 22, 262–269 (2016).

79. Mayer, K.A., Stockl, J., Zlabinger, G.J. & Gualdoni, G.A. Hijacking the Supplies: Metabolism as a Novel Facet of Virus-Host Interaction. Front Immunol 10, 1533 (2019).

80. Thaker, S.K., Ch’ng, J. & Christofk, H.R. Viral hijacking of cellular metabolism. BMC Biol 17, 59 (2019).

81. Cheng, M.L., et al. Metabolic Reprogramming of Host Cells in Response to Enteroviral Infection. Cells 9(2020).

82. Fu, X., et al. Glutamine and glutaminolysis are required for efficient replication of infectious spleen and kidney necrosis virus in Chinese perch brain cells. Oncotarget 8, 2400–2412 (2017).

83. Chambers, J.W., Maguire, T.G. & Alwine, J.C. Glutamine metabolism is essential for human cytomegalovirus infection. J Virol 84, 1867–1873 (2010).

84. Fontaine, K.A., Camarda, R. & Lagunoff, M. Vaccinia virus requires glutamine but not glucose for efficient replication. J Virol 88, 4366–4374 (2014).

85. Li, H., et al. The landscape of cancer cell line metabolism. Nat Med 25, 850–860 (2019).

86. Ray, P., et al. A Pharmacological Interactome between COVID-19 Patient Samples and Human Sensory Neurons Reveals Potential Drivers of Neurogenic Pulmonary Dysfunction. SSRN, 3581446 (2020).

87. Holland, S.J., et al. R428, a selective small molecule inhibitor of Axl kinase, blocks tumor spread and prolongs survival in models of metastatic breast cancer. Cancer Res 70, 1544–1554 (2010).

88. Azulay, R.D., Velloso, M.B., Suguimoto, S.Y., Ishida, C.E. & Pereira Junior, A.C. Acute disseminated paracoccidioidomycosis. Septic shock. Int J Dermatol 27, 510–511 (1988).

89. Cai, H. Sex difference and smoking predisposition in patients with COVID-19. Lancet Respir Med 8, e20 (2020).

90. Dave, N., et al. Functional cooperation between Snail1 and twist in the regulation of ZEB1 expression during epithelial to mesenchymal transition. J Biol Chem 286, 12024–12032 (2011).

91. Zuo, W., Zhao, X. & Chen, Y.-G. SARS Coronavirus and Lung Fibrosis. Molecular Biology of the SARS-Coronavirus, 247–258 (2009).

92. Jarcho, J.A., Ingelfinger, J.R., Hamel, M.B., D’Agostino, R.B. Sr. & Harrington, D.P. Inhibitors of the Renin-Angiotensin-Aldosterone System and Covid-19. N Engl J Med 382, 2462–2464 (2020).

93. Monteil, V., et al. Inhibition of SARS-CoV-2 Infections in Engineered Human Tissues Using Clinical-Grade Soluble Human ACE2. Cell 181, 905–913 e907 (2020).

94. Bouhaddou, M., et al. The Global Phosphorylation Landscape of SARS-CoV-2 Infection. Cell 182, 685–712 e619 (2020).

95. Dowall, S.D., et al. Antiviral Screening of Multiple Compounds against Ebola Virus. Viruses 8(2016).

96. Duclos, G.E., et al. Characterizing smoking-induced transcriptional heterogeneity in the human bronchial epithelium at single-cell resolution. Sci Adv 5, eaaw3413 (2019).

97. Gertz, J., et al. Distinct properties of cell-type-specific and shared transcription factor binding sites. Mol Cell 52, 25–36 (2013).

98. Byers, L.A., et al. Proteomic profiling identifies dysregulated pathways in small cell lung cancer and novel therapeutic targets including PARP1. Cancer Discov 2, 798–811 (2012).

99. Jamieson, A.R., et al. Exploring nonlinear feature space dimension reduction and data representation in breast Cadx with Laplacian eigenmaps and t-SNE. Med Phys 37, 339–351 (2010).

100. Stewart, C.A., et al. Single-cell analyses reveal increased intratumoral heterogeneity after the onset of therapy resistance in small-cell lung cancer. Nature Cancer 1, 423–436 (2020).

