## Supplementary figures and images for "Lung cancer models reveal SARS-CoV-2-induced EMT contributes to COVID-19 pathophysiology"

### Supplemental Figure

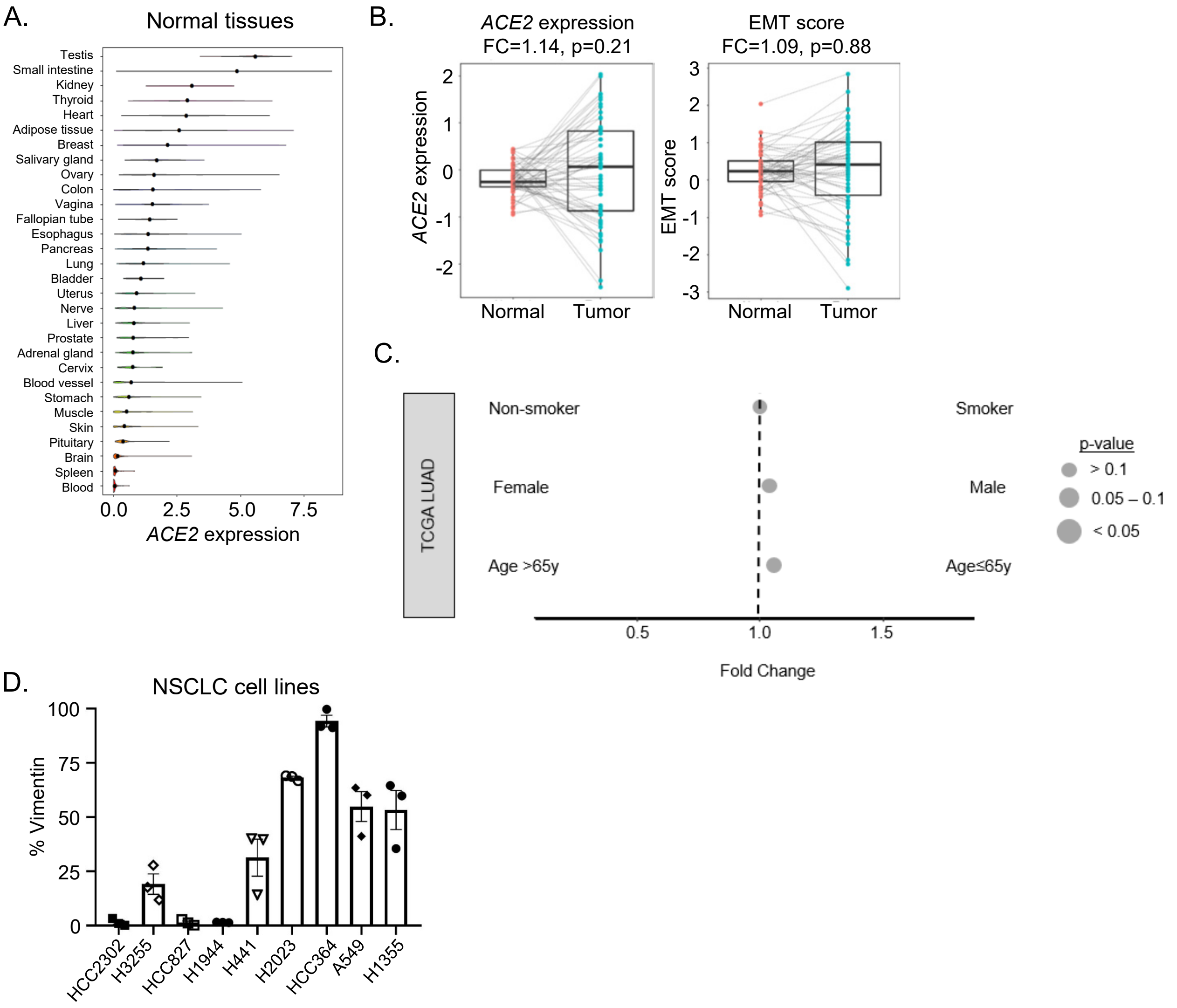


Supplementary Figure 1

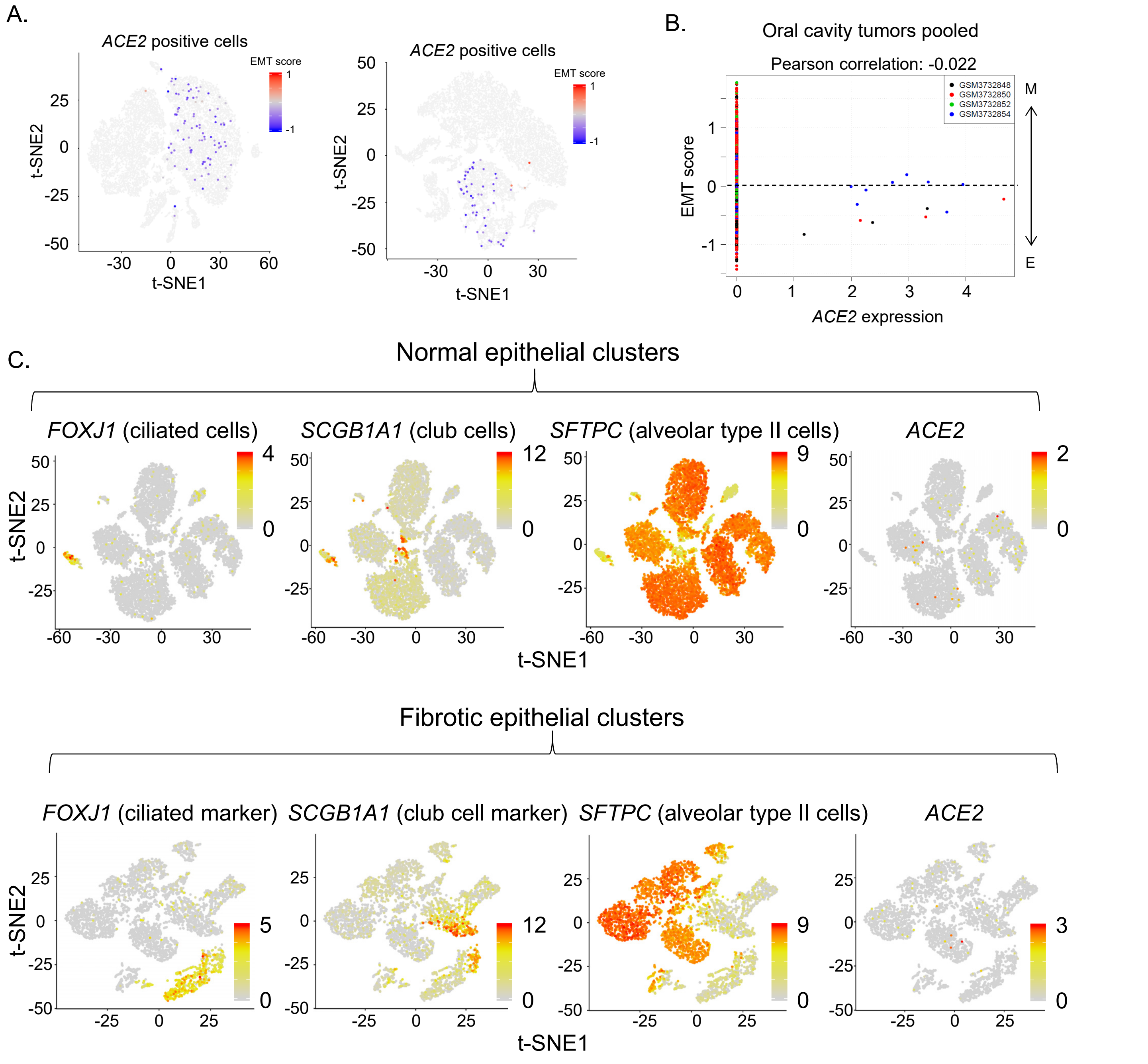


Supplementary Figure 2

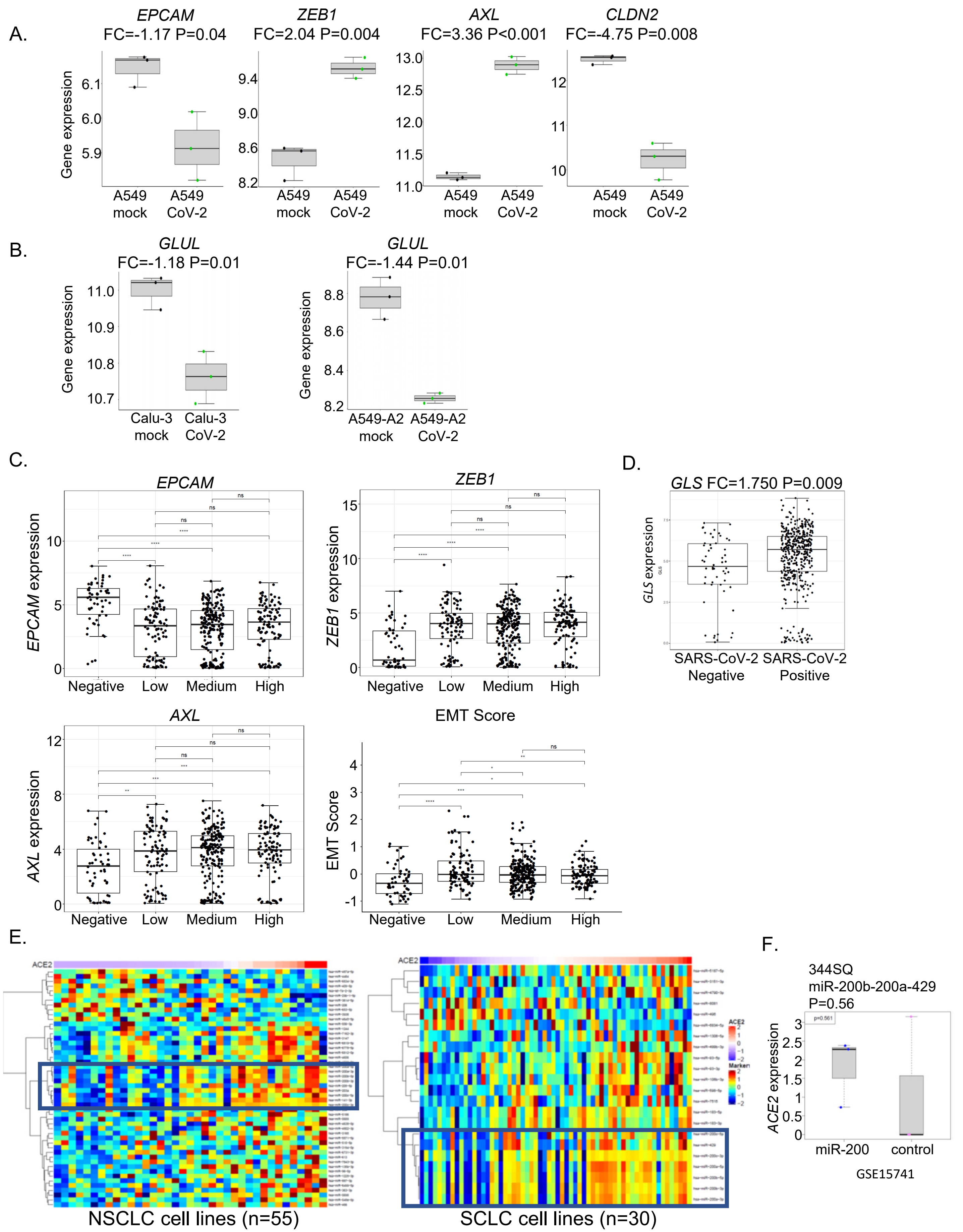


Supplementary Figure 3

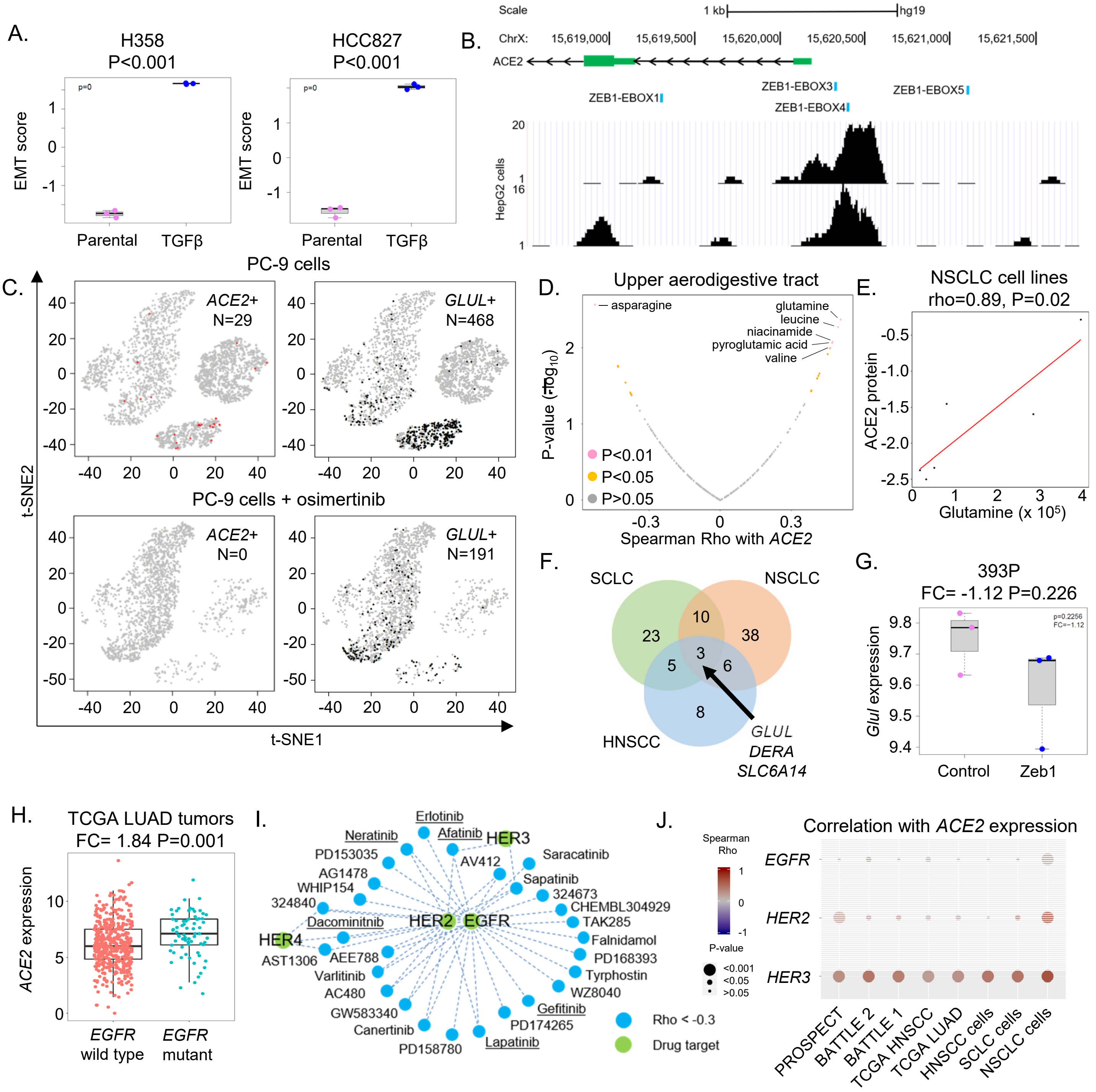


Supplementary Figure 4
