## Supplemental Table 1 for "Lung cancer models reveal SARS-CoV-2-induced EMT contributes to COVID-19 pathophysiology"

| **Dataset** | **Total # of cells/samples** | ***ACE2*+ cells (%)** | ***TMPRSS2*+ cells (%)** | ***POU2F3*+ cells (%)** | ***ACE2+* within *TMPRSS2*+ population (%)** | ***ACE2*+ within *TMPRSS2*- population (%)** | ***ACE2*+ within *POU2F3*+ population (%)** | ***ACE2*+ within *POU2F3*- population (%)** |
| --- | --- | --- | --- | --- | --- | --- | --- | --- |
| Normal lungs | 22,504 cells  (5 lungs) | 110 (0.43%) | 4983 (19.53%) | 270 (1.06%) | 58 cells (1.16%) | 52 cells (0.25%) | 5 cells (2.35%) | 105 cells (0.42%) |
| Fibrotic lungs | 35,610 cells  (8 lungs) | 124 (0.35%) | 4148 (11.65%) | 291 (0.82%) | 59 cells (1.42%) | 65 cells (0.21%) | 6 cells (2.06%) | 118 cells (0.33%) |

Supplemental Table 1. *ACE2* co-expression with *TMPRSS2* and *POU2F3* in normal and fibrotic lungs. *P<0.05, **P<0.01, ***P<0.001.
