## Supplemental Table 2 for "Lung cancer models reveal SARS-CoV-2-induced EMT contributes to COVID-19 pathophysiology"

| **Dataset** | ***ACE2* expression correlated with:** | | | | | | |
| --- | --- | --- | --- | --- | --- | --- | --- |
|  | **EMT score** | ***ZEB1*** | ***ZEB2*** | ***SNAI1*** | ***SNAI2*** | ***TWIST1*** | ***CDH2*** |
| **NSCLC cell lines** | -0.694*** | -0.801*** | -0.477*** | 0.053 | -0.115 | 0.134 | -0.180 |
| **HNSCC cell lines** | -0.408*** | -0.388*** | -0.313** | -0.071 | -0.024 | -0.249* | -0.151 |
| **SCLC cell lines** | -0.401*** | -0.429*** | 0.148 | -0.096 | 0.123 | -0.003 | 0.126 |
| **TCGA LUAD** | -0.286*** | -0.28*** | 0.004 | -0.255*** | -0.175*** | -0.245*** | -0.245*** |
| **TCGA HNSC** | -0.402*** | 0.238*** | -0.214*** | -0.178*** | -0.102* | -0.255*** | -0.210*** |
| **TCGA LUSC** | -0.239*** | -0.070 | -0.093* | -0.230*** | -0.110* | -0.116** | -0.272*** |
| **PROSPECT** | -0.279*** | -0.117* | -0.070 | -0.048 | -0.160** | -0.103 | -0.195*** |
| **BATTLE 1** | -0.367*** | -0.166* | -0.241** | -0.157 | -0.300** | -0.357*** | -0.084 |
| **BATTLE 2** | -0.337*** | -0.196* | -0.211* | -0.344*** | -0.152 | -0.358*** | -0.318*** |

Supplementary Table 2. ACE2 rho correlation values in aerodigestive and respiratory tract cell line and tumor biopsy specimens. *P<0.05, **P<0.01, ***P<0.001.
